# Laminin-α2 is required for the maintenance of the myotendinous junction in vivo

**DOI:** 10.64898/2026.02.18.706289

**Authors:** Julia Schedel, Shuo Lin, Thomas Bock, Dominik Burri, Markus A. Rüegg

**Author notes:** Correspondence to: Markus A. Rüegg.

## Abstract

The myotendinous junction (MTJ) is a critical interface between muscle fibers and tendons, essential for force transmission between muscle and bone. Laminin-α2, a key extracellular matrix (ECM) component, is strongly enriched at this interface. Mutations in the *LAMA2* gene cause LAMA2-related muscular dystrophy (LAMA2 MD), an early-onset severe congenital muscular dystrophy. Here, we examined the MTJ in *dy^W^/dy^W^*mice, a mouse model for LAMA2 MD. We find a strong disruption of MTJ morphology, including altered muscle fiber tips, collagen XXII mislocalization, and reduced muscle tendon interface. As MTJ loading is altered in *dy^W^/dy^W^* mice and MTJ maintenance requires loading and unloading, we also examined MTJ structures upon denervation-induced unloading. While muscle fiber tip morphology resembled that of *dy^W^/dy^W^* mice, collagen XXII distribution was not affected and the muscle–tendon interface was preserved. Finally, proteomic profiling via laser capture microdissection and mass spectrometry revealed significant regional and global shifts in MTJ protein composition in *dy^W^/dy^W^* and denervated mice. Across both models, we identified integrin-associated remodeling as a shared response linked to the perturbed muscle fiber tip morphology. These findings demonstrate that laminin-α2 is required for MTJ stability, and that mechanical unloading contributes to the observed phenotype. Importantly, our results suggest that disruptions in MTJ structure and protein composition may contribute to the pathology observed in LAMA2 MD.

## Introduction

Congenital muscular dystrophies (CMDs) typically present at birth and are associated with significant disability and reduced life expectancy. Mutations in the *LAMA2* gene, which encodes laminin-α2, cause a severe muscular dystrophy characterized by reduced muscle mass and function [1,2]. Laminin-α2 is a particular α chain of the heterotrimeric laminins, which assemble from α, β and γ chains. Laminins are required to form a stable ECM and to connect it to the underlying cell membranes and cytoskeleton by binding to specific muscle cell surface receptors, such as integrins and dystroglycan [1]. Interestingly, the role of laminin-α2 in maintaining structural integrity and tissue function is not limited to skeletal muscle. Indeed, laminin-211 (α2, β1, γ1) is found not only in the endomysial basement membrane (BM) of mature skeletal muscle fibers but also in the heart and the endoneurial BM of peripheral nerves [3]. Peripheral neuropathy, manifested as progressive hind limb paralysis in LAMA2 MD mouse models, highlights the systemic impact of *LAMA2* mutations [4,5]. These findings underscore the complex pathology of LAMA2 MD, revealing the multifaceted roles of laminin-α2 across different tissues.

Laminin-α2 is also enriched at the myotendinous junction (MTJ) [6], a critical anatomical interface between muscle fibers and tendon tissues. The MTJ is essential for force transmission, structural integrity, and coordinated movement. To endure mechanical strain, the MTJ exhibits a highly specialized morphology and molecular composition. It is characterized by interdigitated folds, where tendon tissue protrudes into invaginations of the muscle fiber membrane, enabling robust cell-matrix interactions under mechanical load. At the molecular level, single-nucleus RNA sequencing has shown that MTJ myonuclei are transcriptionally unique compared to body myonuclei [7]. Proteomic studies have highlighted a selective accumulation of ECM proteins at the MTJ [6]. A notable example is Collagen XXII [8], a member of the FACIT (fibril-associated collagens with interrupted triple helix) subgroup. It features a C-terminal collagenous domain that interacts with collagen fibers and N-terminal non-collagenous domains that mediate protein-protein interactions [8]. Knockout of *col22a1* in zebrafish results in loss of MTJ integrity and dysfunctions in force transmissions, underscoring its potential significance, though its precise role remains incompletely understood [9].

To address whether laminin-α2 would also contribute to MTJ structure and function, we investigated the function of laminin-α2 in MTJ integrity using *Lama2*-deficient *dy^W^/dy^W^*mice [10,11]. Morphological and ultrastructural analyses revealed strong disruption of MTJ architecture. To distinguish between a direct effect of laminin-α2 deficiency and the concurrent muscle atrophy and reduced muscle strength, we in addition examined MTJ structure when muscles were denervated for 1 and 4 weeks. Although denervation induced alterations in muscle fiber morphology, it did not recapitulate most of the structural changes observed in *dy^W^/dy^W^*mice. To get some mechanistic insights into these changes, we also examined the MTJ proteome in *dy^W^*/*dy^W^* and denervated muscle, which revealed shared alterations in integrin-associated signaling. These molecular changes indicate that structural disruption at the MTJ is paralleled by adaptive shifts in its signaling pathways. Together, our findings establish laminin-α2 as a pivotal component not only for anchoring the sarcolemma to the ECM but also for maintaining the structural and functional connection between muscle fibers and tendons.

## Results

### Lama2 transcripts and Laminin-α2 are enriched at the myotendinous junction

We analyzed recently obtained single-nuclei RNA sequencing (snRNA-seq) data [12] from *tibialis anterior* muscle to identify, which mononuclear cells and myonuclei subtypes expressed high levels of *Lama2*. Among the mononuclear cells described by Ham et al., 2025, fibroadipogenic progenitors (FAPs), pericytes, muscle satellite/stem cells (MuSCs) and, although with a lower intensity, tenocytes, peri- and endoneurial cells and Schwann cells expressed *Lama2* (Fig. 1A). Interestingly, MTJ and NMJ myonuclei expressed much higher levels of *Lama2* than the body myonuclei (Fig. 1A) and more than 90% of those nuclei expressed detectable levels of *Lama2* (Fig. 1B). To examine the spatial distribution of transcripts, we utilized single molecule RNA fluorescence *in situ* hybridization (sm-FISH) to detect *Lama2* and the MTJ-specific transcript *Col22a1* [8]. In muscle, *Lama2*-positive puncta were detected in specific spots along the laminin-positive muscle BM, while *Col22a1* was not detected (Fig. 1C-E). These results are consistent with the snRNAseq data, which indicated *Lama2* expression in interstitial cells such as pericytes, FAPs and MuSCs (Fig. 1A). At the MTJs, *Lama2* and *Col22a1* expression closely overlapped, marking the specialized transcriptional signature of these nuclei. To test whether local expression of *Lama2* resulted in the accumulation of the protein, we next stained for laminin-α2 protein and visualized the tendon with the antibodies to thrombospondin-4 (Fig. 1F).

**Figure 1:**
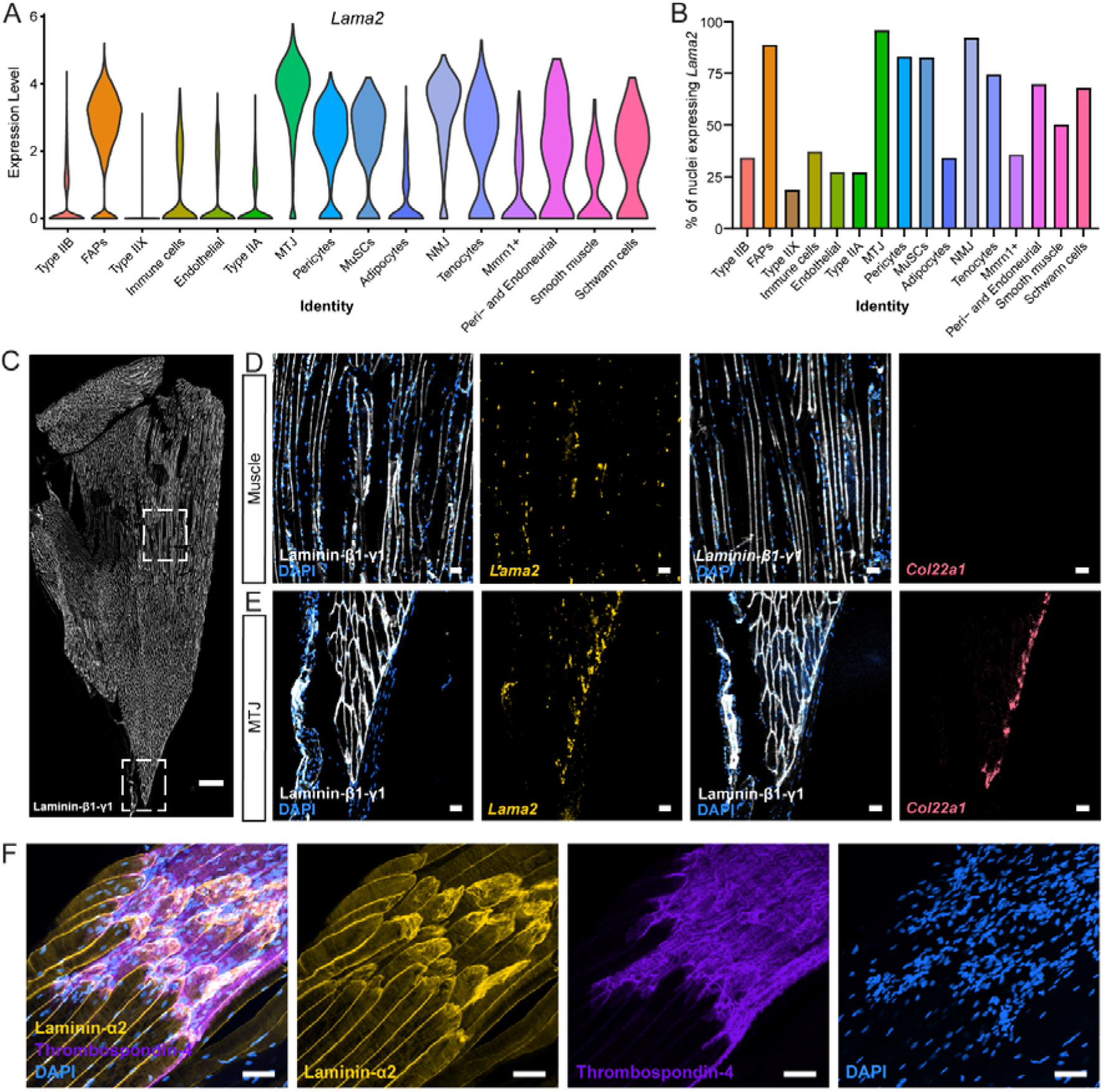
**(A)** Violin plot of *Lama2* expression in the different nuclei populations [12]. **(B)** Bar plot showing the percentage of nuclei expressing *Lama2* within the indicated population. **(C-E)** Representative sm-FISH images for *Lama2* and *Col22a1* in skeletal muscle longitudinal sections, co-stained with antibodies directed against Laminin-β1-γ1. Overview (C) and high-power pictures of muscle fibers (D) and the MTJ region (E). **(F)** Confocal micrographs of muscle-tendon whole-mounts from *extensor digitorum longus* (EDL) from 4-week-old mice. Muscle fibers, stained for laminin-α2 (yellow), insert into the tendon visualized for thrombospondin-4 (lilac). Representative images from n = 3 biological replicates; scale bar = 500 µm (B) and 50 µm (C-E).

Thrombospondin-1, −2 and −4 are known to be enriched at the tendon side of the MTJ [13]. These 3D muscle-tendon whole-mounts confirmed high laminin-α2 abundance at the interface between muscle and tendon (Fig. 1F). Laminin-α2 fully coated the undulating surface of the muscle fiber tip, which appeared as a hilly surface marked by regions of enriched staining. Overall, these data suggest a possible role of laminin-α2 at the MTJ.

### Laminin-α2 deficiency in dy^W^/dy^W^ mice disrupts muscle fiber tip morphology and MTJ ultrastructure

To elucidate the functional significance of laminin-α2 at the MTJ, we utilized the *dy^W^/dy^W^*mouse model. Using 3D whole-mount preparations of *extensor digitorum longus* (EDL), *gastrocnemius* (GAS), and *triceps brachii* (TRI) muscles from 4- to 8-week-old mice, we performed immunostaining with antibodies specific to marker proteins enriched in muscle, MTJ and tendon. In control mice, muscle fiber tips at the MTJ formed rounded, cap-like structures in EDL (Fig. 2A), GAS (Fig. S1A) and TRI (Fig. S1C). In contrast, the tips in all muscles of *dy^W^/dy^W^*mice were rather pointed (Fig. 2B; Fig. S1B, S1D). To get a quantitative measure for the shape of the tips, we determined their circularity. In control mice, the values in all muscles were close to 1, indicating circular muscle fiber tips, while these values dropped significantly in *dy^W^/dy^W^* mice (Fig. 2C). Thus, the absence of laminin-α2 results in pronounced morphological alterations at the MTJ.

**Figure 2:**
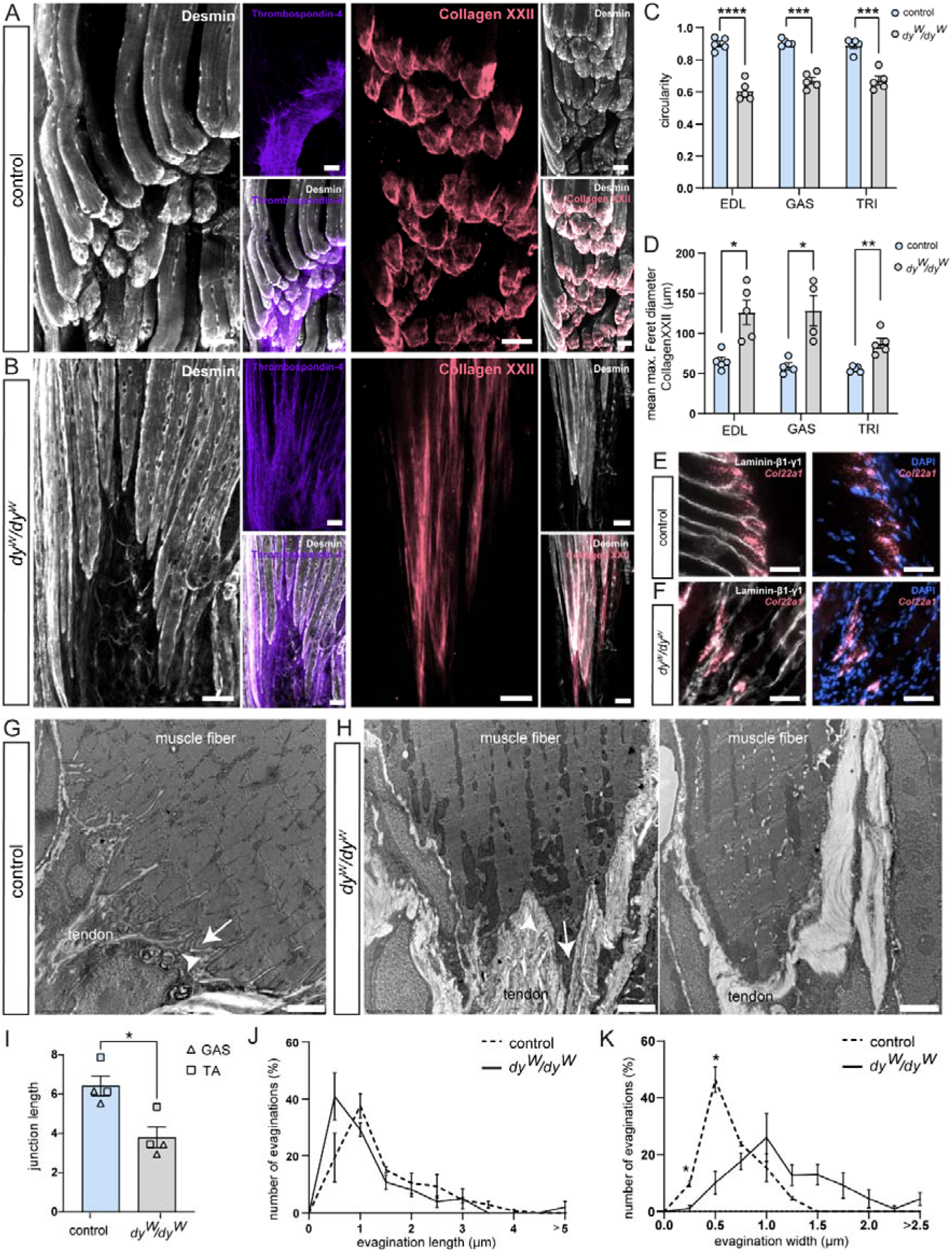
**(A-B)** Confocal micrographs of muscle-tendon whole-mounts from EDL from 4-week-old control and *dy^W^/dy^W^* mice. Muscle fibers are stained for desmin (white), tendons for thrombospondin-4 (lilac). Muscle fiber tips are covered by collagen XXII (red). **(C)** Quantification of muscle fiber tip circularity based on desmin immunostaining in (A) (****P < 0.0001; ***P < 0.001; student t-test; mean ± S.E.M.). Details of the measurements are described in Fig. S3A. **(D)** Quantification of the max. Feret diameter of the collagen XXII signal (**P < 0.01; *P < 0.05; student t-test; mean ± S.E.M.) Details of the measurements are described in Fig. S3B. **(E, F)** Representative images from sm-FISH on longitudinal sections stained for laminin-β1-γ1 showing *Col22a1* localization within control and *dy^W^/dy^W^* MTJ areas. **(G-H)** Transmission-electron micrographs revealing sarcoplasmic invaginations (indicated by arrowheads) and sarcoplasmic evaginations (indicated by arrows) at the muscle-tendon interface of 8-week-old mice. **(I)** Quantification of the junction length from transmission electron micrographs (G, H) (Fig. S3C; *P < 0.05; student t-test; mean ± S.E.M.). **(J-K)** Quantification of the sarcoplasmic evagination length (J) and evagination width (K) (details see Fig. S3D; *P < 0.05; student t-test; mean ± S.E.M.) displayed by the finger-like projections observed in (G-H). Representative images from n = 5 biological replicates for A, B, E, F and n = 3 for G, K; scale bar = 50 µm in A-F and 2 µm in G-H.

The structural difference also resulted in the re-distribution of the MTJ marker collagen XXII. In control mice, collagen XXII was confined to the muscle fiber tip forming a cap-like structure specifically located at the muscle-tendon connection site (Fig. 2A). In *dy^W^/dy^W^*mice, collagen XXII immunoreactivity at the muscle fiber tip was markedly broadened and extended toward the muscle fiber belly (Fig. 2B; Fig. S1B, S1D). This mislocalization was reflected by the significant increase in the maximum (max.) Feret diameter of the collagen XXII signal in all the examined muscles (Fig. 2D). Similarly, *Col22a1* expression was also broadened at *dy^W^/dy^W^*MTJs and extended into the muscle fibers, while the transcripts remained highly concentrated at the MTJ in control mice (Fig. 2E, F). This redistribution of collagen XXII indicates a disruption of its typical spatial organization at the MTJ. Whether the relocalization of collagen XXII represents a compensatory mechanism to mitigate reduced integrity of the MTJ or further compromises the stability of the muscle-tendon interface remains to be elucidated.

To examine the MTJ in the different mice at high resolution, we performed transmission electron microscopy (TEM). In GAS and *tibialis anterior* (TA) of control mice, the MTJ ultrastructure exhibited well-defined interdigitations of the muscle fiber membrane, forming characteristic finger-like processes (Fig. 2G and S2A). Sarcoplasmic evaginations of the muscle extended into the ECM of the tendon, while tendon projections invaded corresponding sarcoplasmic invaginations (Fig. 2G; Fig. S2A). In *dy^W^/dy^W^* mice, muscle fiber endings displayed irregular shapes and a marked reduction in the interdigitation density was apparent at the muscle-tendon interface in both GAS (Fig. 2H) and TA muscle (Fig. S2B). In some cases, the characteristic finger-like processes were entirely absent (Fig. 2H, right image). The junction length, assessed by tracing the finger-like protrusions of the MTJ (Fig. S3C), was significantly reduced in *dy^W^/dy^W^*mice compared to controls (Fig. 2I). We further analyzed the dimensions of the evaginations, specifically measuring their length from the apical point of the ultrastructure to its base, and their width from the left to the right border at the base (Fig. S3D). In TA, sarcoplasmic evagination length was significantly increased compared to control (Fig. S2C, D). In GAS, however, the length of the sarcoplasmic evaginations was not altered but the width was significantly increased (Fig. 2J, K). Thus, while TA shows primarily a change in evagination elongation, GAS exhibits widening, indicating a different response to the loss of laminin-α2 in different muscles. The structural changes indicate a profound disruption of MTJ architecture in *dy^W^/dy^W^* mice.

#### Long-term denervation partially recapitulates morphological alterations observed at the MTJ of dy^W^/dy^W^ mice

*Lama2*-deficient *dy^W^*/*dy^W^* mice are characterized by a progressive muscular dystrophy that manifests already at birth [1,14]. As a consequence, the muscles are much weaker than those of wild-type mice [15]. In addition, *dy^W^*/*dy^W^* mice develop a progressive hindlimb paralysis, caused by the lack of laminin-α2 in the endoneurium of the peripheral nerve [4]. This impairment of nerve conduction further exacerbates muscle weakness and functional decline. As muscle contraction and loading is critical for the maturation of the MTJ [16], we hypothesized that reduced force transmission in *Lama2*-deficient mice may also contribute to the observed MTJ phenotype. To test this, we analyzed the morphology and ultrastructure of the MTJ in 8-week-old mice that were subjected to sciatic nerve transection for one and four weeks.

Following one week of denervation, muscle fiber tips exhibited no morphological changes, as reflected by unchanged muscle fiber tip circularity in GAS (Fig. S4A, S4B) and EDL (Fig. S4C, S4D). In addition, the collagen XXII signal appeared more compressed, concentrating at the muscle fiber tips (Fig. S4B). This compression of the signal resulted in a significantly reduced max. Feret diameter in EDL and GAS (Fig. S4F). By four weeks of denervation, the collagen XXII staining was indistinguishable from the innervated muscle (Fig. 3A, B, D; Fig. S5A, B). As expected, muscle fibers were atrophied and showed a distinct thinning at their fiber tips (Fig. 3B). As a consequence of the muscle atrophy, the tip circularity was reduced in EDL and GAS (Fig. 3C).

**Figure 3:**
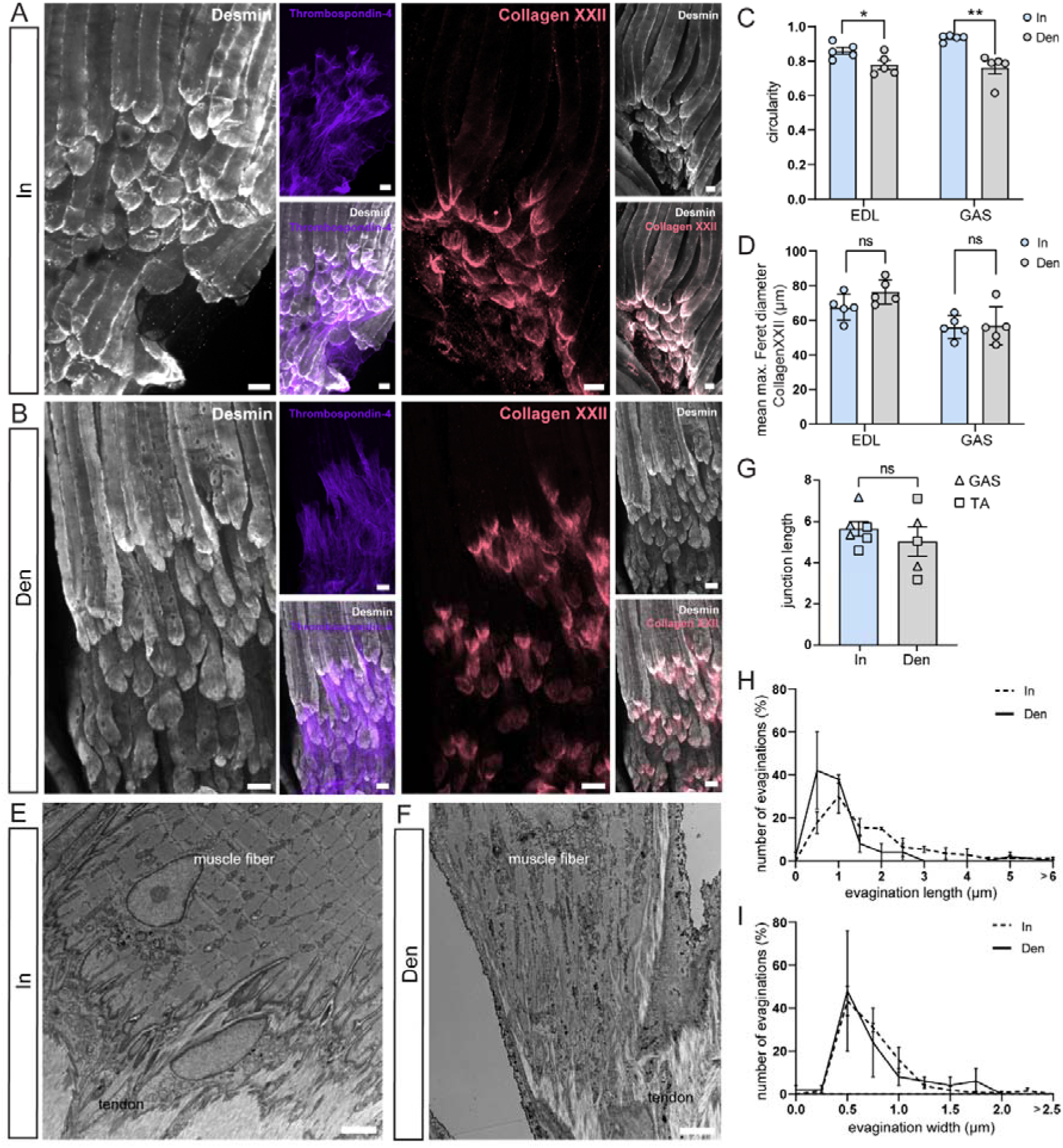
**(A-B)** Confocal micrographs muscle-tendon whole-mounts from *gastrocnemius* (GAS) of 8-week-old mice after 4 weeks of denervation. Immmunohistochemistry for desmin (muscle fibers; white), thrombospondin-4 (tendons; lilac) and collagen XXII (muscle-tendon interface, red). **(C)** Quantification of muscle fiber tip circularity based on desmin immunostaining in (A-B) (**P < 0.01, *P < 0.05; student t-test; mean ± S.E.M.). **(D)** Quantification of the max. Feret diameter of the collagen XXII signal (ns > 0.05; student t-test; mean ± S.E.M.). **(E-F)** Transmission-electron micrographs of the muscle-tendon interface from GAS of innervated (In) and 4-weeks denervated (Den) muscle. **(G)** Quantification of the muscle-tendon interface area from Transmission Electron micrographs in (E-F) (ns > 0.05; student t-test; mean ± S.E.M.). **(H-I)** Quantification of the evagination length (H) and evagination width (I) displayed by the finger-like projections observed in (E-F). Representative images from n = 5 biological replicates for A, B and n = 3 for E-F; scale bar = 50 µm in A-F and 2 µm in G-H.

Despite these alterations, denervation did not affect the junction length of the muscle-tendon interface, as examined by transmission electron microscopy 1 week post-denervation (Fig. S4E, F) or after 4 weeks in GAS (Fig. 3E-G) and TA (Fig. S5C, D). Similarly, evagination length and width remained unaffected after 4 weeks of denervation in GAS (Fig. 3H-I) and TA (Fig. S5E-F). These findings indicate that denervation reduces muscle fiber tip circularity of the MTJ, which is also observed in *dy^W^/dy^W^* mice. However, many of the structural changes are not reproduced by denervation. This highlights that laminin-α2 plays an essential role beyond mechanical force transmission, likely contributing directly to the structural integrity of the MTJ.

#### Proteomic profiling of the MTJ of dy^W^/dy^W^ and 4-weeks denervated mice

To get some insights into the molecular mechanism involved in the changes observed at the MTJ in *dy^W^/dy^W^* mice and upon denervation, we utilized proteomics. We analyzed MTJ and muscle samples from control and *dy^W^/dy^W^* mice from the hindlimb GAS muscle (Table S1) and the forelimb *triceps brachii* muscle (TRI) (Table S2) as the latter is not affected by paralysis. To account for any changes caused by denervation, we also included GAS samples from wild-type mice where one leg was denervated for 4 weeks and the contralateral leg served as an innervated control. Laser capture microdissection (LCM) was used to isolate the MTJ region with as little contamination from adjacent muscle and tendon tissues as possible (Fig. 4A). Proteins were then processed using the single-pot, solid-phase-enhanced sample-preparation (SP3) approach [17] that was optimized for ultra-low input samples, and mass spectrometry was performed using a data-independent acquisition method. This approach ensured a high degree of specificity and sensitivity, enabling a precise characterization of the MTJ proteome. Comprehensive proteomic profiling quantified approximately 6,000 proteins from GAS (Fig. 4B, Table S1) and 3,500 proteins from TRI (Fig. S6A, Table S2). Principle Component Analysis (PCA) revealed distinct segregation of sample types, with separation according to the genotype along PC1 and region (muscle/MTJ) along PC2 (Fig. 4C; Fig. S6B). In the denervated samples, PCA segregated the experimental manipulation along PC1 and again region (muscle/MTJ) along PC2 (Fig. 4D; Fig. S6C). Overall abundance changes on the common set of proteins between the two muscles GAS (x-axis) and TRI (y-axis) within the comparisons LogFC(MTJ control/Muscle control) (Fig. S6D) and LogFC(MTJ *dy^W^/dy^W^*/Muscle *dy^W^/dy^W^*) (Fig. S6E) showed moderate correlations (r(control) = 0.54, r(*dy^W^/dy^W^*) = 0.57) suggesting partial conservation of MTJ-specific protein profiles between muscles.

**Figure 4:**
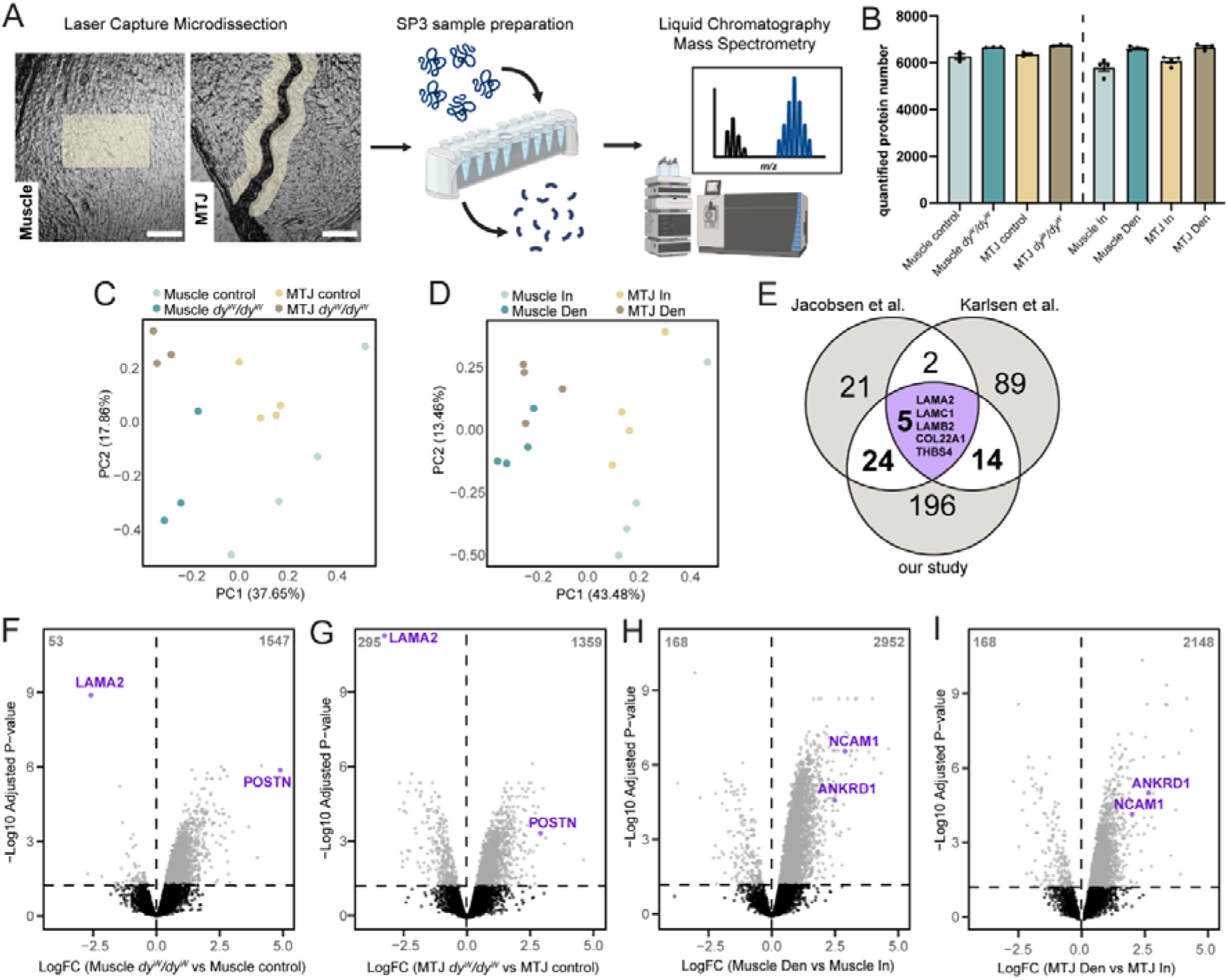
**(A)** Illustration of the proteomics workflow including Laser-capture microdissection of the different sample regions muscle and MTJ from GAS muscle sections of 4 week-old male mice, SP3 sample preparation and LC-MS. **(B)** Bar plot of the number of quantified proteins within each sample type for GAS (Table S1). **(C-D)** Principal component analysis showing the segregation of the samples within each experimental group separately. **(E)** Venn diagram comparing proteins identified as enriched at the MTJ in this study and previous proteomic datasets [6,13]. The dataset from this study includes proteins significantly enriched in wildtype MTJ versus wildtype muscle (P-value < 0.05, LogFC > 1). Protein numbers shown in bold are presented as a heatmap in S6F. **(F-I)** Volcano plots displaying the comparison of *dy^W^/dy^W^* vs control proteome (F-G) and denervated vs innervated proteome (H-I) in the sample region muscle and MTJ separately. The significance level of the adjusted P-value of 0.05 is indicated by a horizontal dashed line.

As quality control, our dataset was compared with previously published datasets in order to assess the specificity of the MTJ proteomic profile [6,13]. Among the three studies, a common core of five proteins (LAMA2, LAMC1, LAMB2, COL22A1, THBS4) was consistently identified as MTJ-enriched, highlighting their robust association with the MTJ across independent experimental approaches (Fig. 4E). Additional overlaps included 14 proteins shared with Karlsen et al. and 24 proteins shared with Jacobsen et al., whereas each dataset also contained distinct proteins not detected in the others (Fig. S6F). The limited overlap likely reflects differences in MTJ isolation and extraction methods used across studies. Together, these comparisons point to a small but conserved molecular core of the MTJ, alongside broader variability between studies.

We identified proteins most significantly altered in *dy^W^/dy^W^* and denervated mice by performing t-test separately for each region. As expected, laminin-α2, was the most significantly downregulated protein in *dy^W^/dy^W^* mice. Additionally, the amount of periostin (POSTN), a marker protein known to be upregulated in muscular dystrophies [18,19], was significantly higher across both analyzed regions in *dy^W^/dy^W^* mice of GAS (Fig. 4F, G) and TRI muscles (Fig. S6G, H). In the comparison between denervated and innervated samples, the known denervation markers Ankyrin repeat domain-containing protein 1 (ANKRD1) and Neural cell adhesion molecule (NCAM1) [20] were significantly upregulated in the denervated condition (Fig. 4H, I), consistent with their established roles in the muscle denervation response. Strikingly, the number of upregulated proteins in each experimental comparison was notably higher than the number of downregulated ones, a trend that has been previously observed in mass spectrometry analyses of 14-day-denervated muscle [20].

#### MTJ-specific protein abundance patterns in dy^W^/dy^W^and denervated mice

To identify distinct patterns of protein regulation across the four experimental conditions, we performed differential expression analysis quantifying interaction and main effect to uncover both region-specific and global changes in protein expression in *dy^W^/dy^W^* and denervated mice (Data S1,2 and Table S3). As muscle and MTJ regions in each sample derived from the same mouse, we used a paired design within the calculation of interaction and main effects. Main effects were used to determine consistent differences in protein abundance between MTJ and muscle across conditions. Interaction effects, in contrast, captured condition-specific differences, identifying proteins that respond differently at the MTJ compared to the muscle in *dy^W^/dy^W^*or denervated mice.

To gain an overview of how MTJ-to-muscle differences behave across conditions, we next compared MTJ-to-muscle ratios between control and disease states using correlation. In *dy^W^/dy^W^* mice, the relationship between MTJ and muscle ratios (LogFC(MTJ *dy^W^/dy^W^*/Muscle *dy^W^/dy^W^*), y-axis) and those of controls (LogFC(MTJ control/Muscle control), x-axis), correlated moderately (r = 0.56, Fig. 5A). In contrast, in the denervated versus innervated samples, MTJ-to-muscle ratios (LogFC(MTJ Den/Muscle Den, x-axis) vs LogFC(MTJ In/Muscle In, y-axis) showed only a weak correlation (r = 0.27, Fig. 5B). In these scatter plots, proteins that fall along the diagonal represent main effects shown in lilac, i.e. their MTJ-to-muscle differences were preserved across conditions. Points above the diagonal indicated stronger MTJ enrichment in *dy^W^/dy^W^* and denervation compared to controls, whereas those below the diagonal represent reduced MTJ enrichment. In contrast, proteins with an interaction effect, highlighted in magenta, deviate from the diagonal and occupy the off-diagonal quadrants, indicating that their regulation differs between MTJ and muscle. Consistent with this, shared main effects between the MTJ and Muscle, shown in lilac, were more pronounced in the *dy^W^/dy^W^* vs control comparison (Fig. 5A). Conversely, interaction-specific effects, highlighted in magenta, were more abundant in the denervated vs innervated comparison (Fig. 5B). These data indicate that in the *dy^W^/dy^W^* model global changes dominate in muscle and MTJ regions whereas region-specific responses predominate in the denervation model. Having established these overall patterns, we next asked how MTJ-enriched proteins behave in *dy^W^/dy^W^* mice, since these candidates are most directly linked to junctional function and pathology. By restricting the analysis to this subset, we could determine which MTJ-associated proteins were consistently altered across regions (main effects) and which showed MTJ-specific regulation (interaction effects).

**Figure 5:**
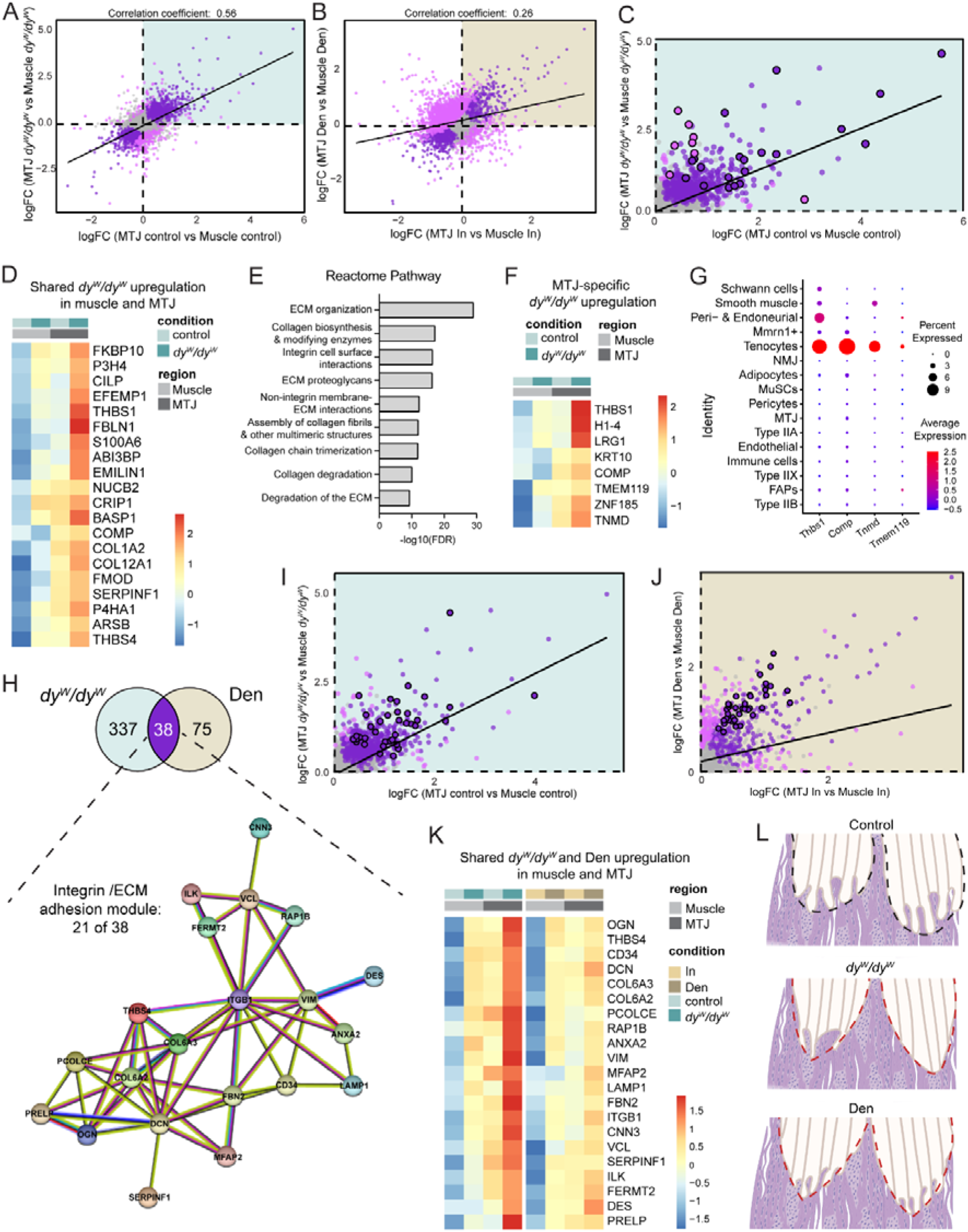
**(A-B)** Pairwise comparisons of regional protein abundance changes between *dy^W^/dy^W^*and control mice (A) and denervated vs innervated mice (B) (Table S1, 3). Main effects (P-value < 0.05) are highlighted with lilac and interaction effects (P-value < 0.05) are highlighted in magenta. Each point represents one protein. A linear solid regression line (black) indicates the trend. The Pearson correlation coefficient is displayed on the plot. **(C)** Zoom into the upper-right quadrant of the pairwise comparison shown in (A). MTJ-enriched proteins (LogFC(MTJ control vs Muscle control) > 0.25), that are upregulated (LogFC(MTJ *dy^W^/dy^W^* vs MTJ control) > 0.25) in the muscle and the MTJ of *dy^W^/dy^W^* mice (P-value(main contrast) ≤ 0.05) are highlighted in lilac with a bold black outline. MTJ-specific proteins (LogFC(MTJ control vs Muscle control) > 0.25) that are upregulated (LogFC(MTJ *dy^W^/dy^W^* vs MTJ control) > 0.25) in the MTJ of *dy^W^/dy^W^* mice (P-value(interaction contrast) ≤ 0.05) are highlighted in magenta with with a bold black outline**. (D)** Heatmap of top 20 most significant MTJ-enriched proteins (LogFC(MTJ control vs Muscle control) > 0.25), that are upregulated (LogFC(MTJ *dy^W^/dy^W^*vs MTJ control) > 0.25) in the muscle and the MTJ of *dy^W^/dy^W^* mice (P-value(main contrast) ≤ 0.05). Tile colors represent scaled protein abundance values. **(E)** Reactome Pathway Analysis of proteins shown in D. **(F)** Heatmap of MTJ-specific proteins (LogFC(MTJ control vs Muscle control) > 0.25) that are upregulated (LogFC(MTJ *dy^W^/dy^W^* vs MTJ control) > 0.25) in the MTJ of *dy^W^/dy^W^* mice (P-value(interaction contrast) ≤ 0.05). Tile colors represent scaled protein abundance values. **(G)** Expression map of marker genes respective for the identified proteins in (F) across nuclear subpopulations [12]. **(H)** Venn diagram displaying the overlap of MTJ-enriched proteins (LogFC(MTJ control vs Muscle control) > 0.25) that display concordant regulation in *dy^W^/dy^W^* and denervated mice (P-value(main effect) < 0.05). A STRING-connected subset [21] comprising 21 of 38 shared proteins formed an integrin/ECM adhesion module. Only integrin-related signaling members are shown. Nodes represent proteins and edges indicate predicted functional associations. **(I-J)** Zoom into the upper-right quadrant of the pairwise comparisons shown in (A) (I) and (B) (J). Proteins in the overlap shown in (H) are highlighted in lilac with a bold black outline. **(K)** Heatmap of the 21 proteins compromising the integrin/ECM adhesion mocule identified in (H) that are upregulated in both *dy^W^/dy^W^*and denervated mice (P-value(main contrast) ≤ 0.05). Color in each tile represents the scaled abundance value. **(L)** Schematic illustration of MTJ morphology in wild-type, *dy^W^/dy^W^* and denervated mice. Dashed lines indicate fiber tip shaping, which is sharpened in *dy^W^/dy^W^*and denervated mice (lilac line) compared to control mice (black line). Created with Biorender.

We first focused on MTJ-enriched proteins with a main effect. In order to select the desired proteins, we applied additional filtering criteria: proteins were required (i) to be enriched at the MTJ under baseline conditions (LogFC(MTJ control vs Muscle control) > 0.25), and (ii) to display a main effect, i.e., to be regulated in the same direction in both MTJ and muscle (P-value(main effect) < 0.05) (Table S3). Within this set, we distinguished up- and downregulation at the MTJ. Applying an upregulation threshold (LogFC(MTJ *dy^W^/dy^W^* vs MTJ control) > 0.25) identified a subset of MTJ-specific proteins with increased abundance in *dy^W^/dy^W^* mice (Fig. 5C-D), whereas no proteins met equivalent downregulation criteria. This showed that the MTJ response in *dy^W^/dy^W^*was dominated by elevated expression rather than loss of MTJ-enriched components. The expression patterns of the upregulated candidates across conditions, confirmed their MTJ enrichment in *dy^W^/dy^W^* mice (Fig. 5C, D). Increased abundance of proteins such as Fibulin-1 in the muscle and at the MTJ in *dy^W^/dy^W^*mice was further confirmed using immunostaining (Fig. S7A, B). Having identified a subset of MTJ-enriched proteins consistently increased in *dy^W^/dy^W^*mice, we next asked which biological processes they are associated with. Pathway analysis of these candidates revealed enrichment for ECM remodeling and organization (Fig. 5E), indicative of fibrosis. This finding is consistent with the known pathology of LAMA2-MD, where excessive ECM deposition and fibrosis are hallmarks of disease progression [1,15].

Finally, we focused on proteins showing an interaction effect. These represent cases where disease-induced changes are specific to the MTJ and not mirrored in muscle, highlighting distinct molecular responses at the junction. To identify such proteins, we applied filtering criteria: (i) baseline MTJ enrichment (LogFC(MTJ control vs Muscle control) > 0.25), and (ii) differential regulation at the MTJ in *dy^W^/dy^W^* relative to control mice (P-value(interaction effect) < 0.05) (Table S3). Candidates were then classified as increased (LogFC(MTJ *dy^W^/dy^W^*vs MTJ control) > 0.25) or decreased (LogFC(MTJ *dy^W^/dy^W^* vs MTJ control) < –0.25) ). No proteins fulfilled the criteria for decreased abundance, which is consistent with our main-effect analysis. The elevated proteins were significantly increased at the MTJ in *dy^W^/dy^W^* mice (Fig. 5F). While many also showed a tendency toward higher abundance in muscle that did not reach statistical significance, two proteins—Keratin 10 (K10) and Cartilage Oligomeric Matrix Protein (COMP)—exhibited an opposite trend with reduced levels in muscle.

Interestingly, many of the MTJ-specific proteins identified in this analysis (Fig. 5F) were classically associated with tendon and cartilage development, raising the possibility that their upregulation reflects altered cellular composition at the junction. As a complementary approach, we therefore interrogated snRNAseq data to assess whether these proteins correspond to known tenocyte markers. Transmembrane Protein 119 (TMEM119), Tenomodulin (TNMD), COMP, and Thrombospondin-1 (THBS1) are specifically expressed in tenocytes (Fig. 5G). sm-FISH confirmed MTJ-specific enrichment of *Tnmd* in *dy^W^/dy^W^*mice, together with *Col22a1* (Fig. S7C, D). These findings suggested an infiltration or expansion of tenocytes at the MTJ in *dy^W^/dy^W^* mice. To determine whether the increased tenocyte number resulted from tendon involvement in LAMA2 MD, we performed LC-MS on dissected muscle and tendon samples (Fig. S7E, Table S4). Pairwise t-tests revealed over 3,000 significantly altered proteins (adj. P-value (*dy^W^/dy^W^* vs control) < 0.05) in the muscle region (Fig. S7F) compared to only 133 in the tendon region (Fig. S7G). Notably, none of the MTJ-specific proteins identified through interaction analysis was altered in tendon, indicating that these changes were localized to the muscle rather than arising from the tendon proteome. Further work will be required to clarify whether the tenocyte response detected at the MTJ exacerbates junctional pathology or represents a compensatory attempt at repair.

Because both *dy^W^/dy^W^* and denervation models exhibited shared structural changes like reduced muscle fiber tip circularity, we next asked whether they show common molecular changes that might contribute to this phenotype. To address this, we applied filtering criteria to focus on MTJ-enriched proteins that were consistently altered across regions in both models. Specifically, we considered proteins that were (i) MTJ-enriched under baseline conditions (LogFC(MTJ control vs Muscle control) > 0.25) and (ii) displaying a main effect, meaning they were regulated in the same direction in both MTJ and muscle (P-value(main effect) < 0.05) (Table S3). In line with our previous analyses, we restricted the comparison to upregulated proteins, as no downregulated candidates were detected within the main-effect contrasts in *dy^W^/dy^W^* mice. Comparison of the filtered protein sets from *dy^W^/dy^W^* and denervation revealed 38 overlapping candidates out of 450 (11.8%) (Fig. 5H-K) indicating a substantial shared molecular response. To investigate functional relationships within this overlap, STRING network analysis identified an integrin-centered interaction network compromising 21 of the 38 shared proteins, enriched for extracellular matrix, cytoskeletal, and adhesome-associated proteins. ITGB1 occupied a central position within the network, forming extensive connections with ILK, FERMT2, VIM, SPARC, and multiple collagen and proteoglycan components, consistent with its role as a major signaling and structural node (Fig. 5H). Given the established role of integrins in mechanosensing and force transmission, these data suggest that altered integrin signaling contributes to the muscle fiber tip and MTJ morphological changes observed in both *dy^W^/dy^W^* and denervated muscle (Fig. 5L). Immunostaining for integrin β1d confirmed elevated total protein abundance in muscle fibers and at the MTJ in *dy^W^/dy^W^* and denervated mice (Fig. S7H–K). In contrast, immunostaining for the activated conformation of β1 integrin showed only a modest increase (Fig. S7L-O), indicating that integrin activation does not scale proportionally with total protein levels. This disparity indicates that increased abundance of integrin pathway components is not accompanied by a proportional increase in force-dependent integrin activation, consistent with the reduced mechanical loading present in both *dy^W^/dy^W^*and denervated muscle. Immunostaining further validated the upregulation of an additional ECM-associated protein, Fibulin-2, in both muscle and MTJ regions (Fig. S7P–S).

## Discussion

The present study provides novel insights into the role of laminin-α2 in organizing the muscle–tendon junction. By combining *Lama2*-deficient and denervated mouse models with ultrastructural, transcriptomic, and proteomic approaches, we show that laminin-α2 is indispensable for MTJ maturation, structural stability, and ECM organization.

Laminin-α2 is highly enriched at the MTJ, as shown by our snRNA-seq, sm-FISH, and 3D immunohistochemistry (Fig. 1A-F), in agreement with human MTJ proteomics [6]. This prominent localization likely explains why its absence in *dy^W^/dy^W^*mice caused striking alterations in fiber tip morphology and MTJ ultrastructure. Muscle fiber tips lost their circularity and adopted pointed shapes (Fig. 2A-C; Fig. S1A-D), resembling the embryonic phenotypes described for collagen XXII, where deposition progresses from aligned at E13.5 to jagged at E18.5 and highly interdigitated at P21 [16]. In *dy^W^/dy^W^*mice, however, collagen XXII remained extended into the fiber belly up to early adulthood (Fig. 2A-D; Fig. S1A-D), indicating a failure of developmental progression. Transmission electron microscopy confirmed profound structural disruption, with a marked reduction in both the width (Fig. 2K) and number (Fig. 2I) of sarcoplasmic evaginations. The resulting loss of interdigitations decreases the attachment surface, concentrates stress at the interface, and increases susceptibility to fiber detachment and contraction-induced damage. Such chronic instability is consistent with the progression in LAMA2 MD, where repeated injury drives cycles of degeneration, regeneration, and fibrosis [1]. These ultrastructural disruptions likely stem from the unique molecular properties of laminin-α2. By self-polymerizing and anchoring to cell surface receptors, laminin-α2 establishes a scaffold for provisional ECM assembly. Its interaction with nidogen-1 further links laminins to type IV collagen, generating a co-polymerized network that connects the sarcolemma to the matrix [22,23]. Loss of this anchoring system prevents proper MTJ maturation and compromises mechanical stability. Notably, the structural remodeling observed in *dy^W^/dy^W^* mice parallels findings in mdx mice, where both the width and length of evaginations are reduced, diminishing the muscle-tendon interface [24]. Similar reductions have been reported in very old rats [25], which also show impaired lateral force transmission. Together, these observations suggest that reduced interdigitation is a shared mechanism of mechanical instability across dystrophies and aging.

Denervation experiments further revealed how mechanical loading shapes MTJ organization. Reduced contractility reproduced the pointed fiber tips seen in *dy^W^/dy^W^* mice (Figs. 3A-C), suggesting that the absence of contractile forces alters MTJ structure. Similar observations were made by Lipp et al. [13] in a murine model of muscular dysgenesis (mdg) in which a mutation in a calcium ion channel (CACNA1S) disrupts muscle excitation-contraction coupling [26,27]. In our denervation experiments, ECM deposition of proteins, such as collagen XXII, was reduced; yet, unlike *Lama2* deficiency, denervation did not induce collagen XXII elongation (Fig. 3A, B, D) or a reduction in junction length (Fig. 3G). These results indicate that laminin-α2 is uniquely required for spatial ECM organization.

Consistent with this, proteomic analysis revealed that laminin-α2 levels at the MTJ remain unchanged upon denervation, suggesting that the preserved presence of laminin-α2 helps maintain overall MTJ architecture and may explain why structural alterations in denervated muscle differ from those observed in *dy^W^/dy^W^* mice. More severe models of mechanical unloading, such as immobilization or hindlimb suspension [28,29] have been shown to cause a reduced muscle-tendon interface [28,29]. In contrast, sciatic nerve transection does not completely abolish passive mechanical loading, as the paralyzed limb is still dragged. Hindlimb suspension, however, eliminates both active and passive loading and therefore likely imposes a greater reduction in mechanical strain. This difference may explain why certain ultrastructural alterations observed in the hindlimb-suspended rats were not recapitulated following denervation alone.

These comparisons suggest that the molecular pathways sensing and transmitting mechanical cues may also contribute to the MTJ phenotype in LAMA2 MD mice as both converged on a common phenotype of muscle fiber tip thinning (Fig. 5L). Proteomic analysis identified shared upregulation of integrin-associated proteins, with ITGB1 emerging as a central hub (Fig, 5H, K). This upregulation was detectable in muscle tissue but was even more pronounced at the MTJ, suggesting a localized response at the site of highest mechanical demand. Notably, several proteins upregulated in both models, including vinculin (VCL), Kindlin-2 (FERMT2), and RAS-like small GTP-binding protein 1b (RAP1B), are part of the canonical FAK-associated adhesome [30,31], and integrin-linked kinase (ILK) and Parvin (PARVB) form the ILK–PINCH–Parvin complex, a parallel integrin effector module that anchors actin and cooperates with FAK-dependent mechanotransduction [32,33]. Similar compensatory increases in integrin pathway members have been reported in mdx mice, where α7 integrin, paxillin, and FAK are upregulated despite weakened ECM–cytoskeleton attachment [34,35].

Moreover, muscle-specific deletion of ILK causes a severe dystrophic phenotype with marked disruption of myofascial and myotendinous junctions, displacement of focal adhesion proteins (vinculin, paxillin, FAK), and fiber tip irregularities [36]—closely mirroring defects seen in α7-integrin deficiency.

Under physiological conditions, mechanical loading activates β1-integrin signaling and downstream effectors such as focal adhesion kinase (FAK), whereas unloading models, such as hindlimb suspension, disrupt focal adhesions and attenuate this pathway [37,38]. In this context, the integrin pathway upregulation observed in our datasets is unlikely to reflect enhanced mechanotransductive signaling. Instead, although total β1 integrin abundance is increased (Fig. S7H-K), the activated conformation of β1 integrin is only modestly upregulated (Fig. S7L-O), indicating that integrin activation does not scale proportionally with protein abundance. This dissociation is consistent with reduced mechanical loading in both *dy^W^/dy^W^* and denervated muscle and suggests compensatory remodeling or accumulation of adhesion components rather than productive force-dependent signaling. Taken together, the integrin-associated remodeling shared between *dy^W^/dy^W^* and denervated MTJs likely represents an adaptive response to structural vulnerability, attempting to stabilize the reduced anchoring surface and maintain MTJ integrity.

Beyond structural disruption, laminin-α2 deficiency induced broad ECM remodeling. Proteomics revealed enrichment of fibrosis-associated proteins like thrombospondins, FMOD and collagens as shared alteration between muscle and MTJ (Fig. 5C, D). This aligns with previous reports that in laminin-α2-deficient mice there is excessive accumulation of ECM [39,40]. Pathway analysis indicated concurrent activation of ECM deposition and degradation (Fig. 5E), suggesting that fibrosis results from disrupted turnover rather than unidirectional accumulation. Similar principles have been observed in collagen XII deficiency, where increased muscle elasticity altered matrix properties and induced a form of functional unloading [41]. In dystrophic conditions, reinforcement of connective tissue may thus help to withstand mechanical stress [42]. This mechanism is particularly relevant for the MTJ, which experiences high loading forces.

At the MTJ specifically, we detected upregulation of tendon- and cartilage-associated proteins such as TNMD, TMEM119, COMP (Fig. 5C, F, S7C, D). This finding is consistent with reports of increased telocyte numbers after atrophy [43] and exercise [44,45], supporting the view that cells accumulate at the MTJ in response to conditions acquiring adaptation. Telocytes are a specialized type of interstitial cell characterized by very long, thin cellular extensions that allow them to form networks and engage in paracrine signaling with neighboring cells [46]. While telocytes have been described in this context, our proteomic and transcriptomic data particularly highlight tenocytes as key contributors. Tenocytes, in particular, are highly mechanosensitive, capable of modulating ECM composition in both anabolic and catabolic directions [47]. Mechanical loading promotes collagen and proteoglycan synthesis, whereas overload triggers pro-inflammatory signaling and MMP-mediated degradation [48,49]. These dual functions suggest that tenocytes could either mitigate or exacerbate MTJ pathology, depending on the mechanical context. Interestingly, despite the presence of tenocyte markers at the MTJ, analysis of isolated muscle and tendon samples revealed minimal alterations in the tendon proteome of *dy^W^/dy^W^* mice (Fig. S7G). This indicates that molecular remodeling likely occurs locally at the MTJ and might be primarily driven by muscle pathology rather than intrinsic tendon dysfunction. Taken together, these findings indicate that the MTJ may serve as a distinct adaptive niche where fibrosis and cell-mediated ECM remodeling act not only as hallmarks of pathology but perhaps also as compensatory mechanisms to stabilize the highly stressed muscle–tendon interface.

In summary, our study highlights the essential role of laminin-α2 in maintaining the structural and molecular integrity of the MTJ. The loss of laminin-α2 leads to disrupted MTJ architecture, altered ECM composition, and increased fibrosis, contributing to the pathological features of LAMA2 MD. Importantly, our findings point to dysregulated integrin signaling, as a shared mechanism underlying the morphological changes observed in both laminin-α2 deficiency and denervation. This compensatory upregulation of adhesion complexes likely represents an attempt to counteract fiber tip thinning and stabilize the MTJ. Further studies are needed to determine whether modulation of integrin pathways can be leveraged therapeutically in diseases associated with laminin-α2 deficiency.

## Experimental procedures

### Mice

As mouse models for LAMA2 MD, we used *dy^W^/dy^W^*mice ([50], [10]) [B6.129S1(Cg)-Lama2tm1Eeng/J; available from the Jackson Laboratory stock #013786]. Control mice were either wild-type or heterozygous. Female and male mice aged 4 – 8 weeks were used for the experiments. To facilitate access to water and food, cages dystrophic mice were equipped with long-necked water bottles and supplied with wet food from weaning onward. All mouse experiments adhered to federal guidelines for animal experimentation and were approved by the authorities of the Canton of Basel-Stadt (Schlachthofstrasse 55, 4056 Basel, Switzerland).

### Genotyping

Mice were PCR genotyped using the following primers: Lama2: FWD: TGCCCTTTCTCACCCACCCTT, REV: GTTGATGCGCTTGGGAC, lacZ insert 5’end: CGACAGTATCGGCCTCAG. 246 bp, 436 bp and both products (436 and 246 bp) represent homozygous, knock-out and heterozygous presence of PCR products, respectively.

### Denervation

Mice were anesthetized by inhalation of isoflurane. A small skin incision was made between the sciatic notch and the knee, and the sciatic nerve was exposed by gentle separation of the underlying muscles. The nerve was elevated using a glass hook and transected by removing a 5=mm segment.

The open wound was closed by surgical clips and mice were returned to their cages. Buprenorphine (0.1=mg/kg body weight) was administered 1=h before surgery and for 2 d postoperatively.

### Wholemount staining

In brief, dissected muscle-tendon units were washed in 1× PBS and fixed in 4% PFA for 20 min at room temperature (RT). Samples were then washed three times in 1× PBS for 10 min each. Muscles were placed in a 24-well plate, blocked, and permeabilized for 3 h in blocking solution (1× PBS, 5% donkey serum, 0.5% Triton-X) under constant shaking at RT. For experiments using mouse primary antibodies, mouse-on-mouse reagent (Vector Laboratories) was included in the blocking solution at a 1:40 dilution. Primary antibodies were diluted in the blocking solution and incubated over night at 4°C. After three PBS washes, samples were incubated with secondary antibody diluted in blocking solution for 1 h at RT. A complete list of antibodies is provided in Supplementary Data Antibodies and Probes (Table S5). Following four additional PBS washes, samples were mounted using ProLong™ Gold Antifade (Invitrogen). Imaging was performed using a SpinD confocal microscope (Olympus). Images are representative of n = 5 biological replicates.

### Single-molecule fluorescent in situ hybridization (sm-FISH)

For RNA detection on muscle cross-sections, we used the RNAscope V2 kit (ACD Bio), following the manufacturer’s protocol. Briefly, fresh-frozen 10 µm muscle cross-sections were fixed in 4% PFA for 15 minutes on ice, then dehydrated through an ethanol series (50%, 70%, and 100%) at room temperature. Tissue digestion was performed using protease IV for 30 minutes. After PBS washes, cross-sections were placed in a humid chamber with hybridization probes, followed by several amplification steps. A complete list of probes is provided in Supplementary Data Antibodies and Probes (Table S5). Sections were then stained with DAPI and mounted using ProLong™ Gold Antifade (Invitrogen). Images were acquired using a Zeiss LSM 700 confocal microscope.

### Electron microscopy

Mice were perfused with fixative (2.5% glutaraldehyde, 2% PFA in 0.1 M sodium cacodylate, pH 7.4) and post-fixed in the same solution for 2 h at 4°C. After three washes with cacodylate buffer 1 (0.1 M, pH 7.2), the samples were incubated in freshly prepared cacodylate buffer 2 (reduced osmium in cacodylate buffer, pH 7.2) for 1 h at RT. Samples were then rinsed several times in cacodylate buffer 2 and transferred to freshly prepared 1% tannic acid in cacodylate buffer (pH 7.2) for 1 h at 4°C. After thorough washing with ddH_2_O, samples were stained with 1% aqueous Uranyl Acetate for 1 h at 4°C in the dark.

Specimens were dehydrated in an ethanol series (30, 50, 75, 95% and 100%) and, after three changes of 100% ethanol, samples are washed in propylene oxide and finally infiltrated in graded series of Epon 812 Hard resin (Electron Microscopy Science) in propylene oxide at RT. After placing of the samples in freshly made 100% Epon 812 Hard resin, embedding was carried out in a 60°C oven for 48 h until complete hardening of the resin based on the protocol described by Strasborg et al. [51].

Semi-thin longitudinal sections were cut with a glass knife, and ultrathin sections (70 nm) were prepared using a diamond knife, placed on copper grids and coated with Parlodion film and a carbon layer. Sections were contrasted with uranyl acetate and lead citrate. The samples were observed in a FEI Tecnai G2 Spirit Transmission Electron Microscope (Thermo Fisher Scientific) operated at 80 kV and images were acquired with an EMSIS Veleta camera using the RADIUS software (EMSIS). Images are representative of n = 3 biological replicates.

### Image processing

Image analysis in case of the immunostaining on muscle-tendon whole-mounts was performed on maximum intensity projections of the z-stacks using I_MAGE_J software. To describe the morphology of muscle fiber tips, the circularity of the fibers was measured in FIJI. For measurement, the tool ‘elliptical selection’ was fitted (Fig. S3A) in the outline of a muscle fiber tip and measured using the variable circularity. A perfect circle has a circularity of 1 and a line a value of 0. Circularity was calculated using the formula: circularity = 4π(Area/(Perimeter^2^)).

The measurement of the expansion of Collagen XXII at individual muscle fiber tips can be divided into 6 steps (Fig. S3B; steps 1-6). First, a raw confocal image was converted to a composite images (step 1), only the Collagen XXII channel was selected and z-projected (step 2). A threshold was set based on the staining intensity (values ranging from 1 to 2), and a binary image was created (step 3). The Collagen XXII area of one muscle fiber was outlined using the ‘Polygon selection’ tool. From this a mask was created (step 4). The mask was inverted (step 5), and the binary image was subtracted from the mask using the ‘Image Calculator’ tool (step 6). Finally, the maximum Feret diameter of Collagen XXII areas was measured.

To quantify the muscle-tendon interface we used the ’Polygon Selection’ tool to surround the finger-like projections of the MTJ (Fig. S3C). This tracing provided the perimeter, representing the length of the interface. To normalize the measurement, the perimeter was divided by the diameter, yielding a standardized value for the junction length of the muscle-tendon interface.

Transmission electron micrographs were analyzed using ImageJ software. For each image, the scale was manually set according to the included 2 µm scale bar. To facilitate evagination measurements, the ’Rectangular Selection’ tool was used to draw a region of interest (ROI) around each individual ultrastructure (Fig. S3D). The apical point and the basement of the ultrastructure defined the rectangle’s boundaries. The evagination length was measured as the longer side of the rectangle, extending from the tip of the ultrastructure to the end facing the interior of the muscle fiber. The evagination width was defined as the shorter side of the rectangle, spanning from the left to the right border of the ultrastructure.

### Protein extraction and sample preparation for MS-based proteome analysis

TMT experiment:

Muscle and tendon samples were prepared for global proteomics analysis using tandem mass tags (TMT) as described previously [52] with a few modifications. In brief, samples were dissected using scissors from gastrocnemius (GAS) muscle-tendon group. Samples were lysed using a PIXUL Multi-Sample Sonicator (Pulse = 50, PRF = 1, Process Time = 20 min, Burst Rate = 20 Hz). 50 µg of total protein was processed using S-Trap™ micro spin columns (Protifi) following the vendor’s instructions. Peptide aliquots (5 μg) were labelled with TMTpro 18-plex isobaric tandem mass tags (Thermo Fisher Scientific), fractionated by high-pH reversed-phase separation into 12 fractions and dried.

Laser Microdissection Experiments:

Laser capture microdissection (LCM) was performed using a PALM MicroBeam system (Zeiss). MTJ areas were identified by brightfield microscopy using a 20x objective lens. Approximately 0.5 mm² of MTJ and muscle tissue was excised and collected by laser-pressure catapulting into low-adherent PCR tubes (Zeiss) containing 10 µL of SDS lysis buffer (5% SDS, 10 mM TCEP, 100 mM TEAB). Approximately 15 MTJ areas were captured per biological replicate, resulting in 32 samples for gastrocnemius and 16 samples for triceps.

Microdissected samples were processed using the single-pot, solid-phase-enhanced sample-preparation (SP3) approach as described [17] with the following modification: Proteins were digested in a 10 µL digestion mix (0.02% DDM, 100 mM TEAB, 10 ng/µL trypsin) at 37°C for 12 h, collected and dried.

### Mass Spectrometry Analysis

All mass spectrometry proteomics data associated with this manuscript have been deposited to the ProteomicsXchange consortium via MassIVE (https://massive.ucsd.edu) with the accession number MSV000096882 / PRIDE PXD059914.

TMT experiment:

250 ng of TMT labeled peptides were LC-MS analyzed using an Orbitrap Eclipse Tribrid Mass Spectrometer coupled to an Ultimate 3000 nano system and a FAIMS Pro interface (Thermo Fisher Scientific) as described [52].

Laser Microdissection Experiments:

Dried peptides were resuspended in 0.1% aqueous formic acid, 0.02% DDM (n-Dodecyl-B-D-maltoside) and analyzed by LC–MS/MS using a timsTOF Ultra Mass Spectrometer (Bruker) and a Vanquish Neo (Thermo Fisher Scientific) or a Orbitrap Eclipse Tribrid Mass Spectrometer coupled to an Ultimate 3000 nanoLC system and a FAIMS Pro interface (all Thermo Fisher Scientific). The TimsTOF Ultra was operated in dia-PASEF mode and MS1 and MS2 scans were acquired over a mass range from 100 to 1700 m/z with ion mobility separation. Concerning the Orbitrap Eclipse Tribrid data was acquired in DIA mode with the FAIMS compensation voltage set to −45 V. MS1 spectra were acquired at a resolution of 60,000 and a scan range of 400 to 1000 m/z.

### Protein Identification and Quantification

TMT experiment:

Raw-files were searched against a Mus musculus SwissProt database (download Jan 2^nd^, 2024) using the SpectroMine software (Biognosys) and further statistically analyzed using our in-house generated SafeQuant R script (v2.3.4) [53] as described [52]. The final processed data table is provided in Table S4.

Laser microdissection experiments:

The acquired files were searched using the Spectronaut directDIA workflow (v19, Biognosys) against the same database as above. Quantitative data were analyzed using the MSstats R package (v.4.13.0) with default normalisation and AFT model-based imputation as described [54]. Log2 intensities were loaded into R (v4.4.1). PCA allowing for missing values was obtained with pcaMethods::pca v1.96.0 using the non-linear iterative partial least squares algorithm (method=”nipals”) on the centered data (center=TRUE and scale=”none”). The object was saved as SingleCellExperiment v1.26.0 object for interactive exploration in a Shiny application with iSEE v2.16.0 (Data S1). Outlier samples were identified based on their expression levels of LAMA2 and NCAM1, as well as their segregation within the Principal Component Analysis (PCA) analysis. The final processed data tables are provided in Tables S1 and S2.

### Differential expression analysis

Differential expression on the paired samples was performed separately for *dy^W^/dy^W^* vs control and Den vs In. A custom design matrix was created using the mouse number as primary factor and adding the condition and region factors manually (Data S2). Linear model was fit with limma::lmFit v3.60.6 with default method least squares. Contrasts were manually defined, fitted with limma::contrastsFit and statistics were moderated with empirical Bayes using limma::eBayes. P-values were adjusted for multiple testing with Benjamini & Hochberg method. Note: missing values are handled by limma. PCA was performed using the prcomp function in R on a scaled and centered matrix of protein abundances. PCA, volcano plots as well as scatter plots were generated using ggplot2. For correlation analysis, proteins with missing values in either comparison were removed to ensure accurate calculations. Pearson’s correlation coefficient was calculated using the cor function in R. Heatmaps were generated using the mean Log2 intensities across replicates, with missing values replaced by zero. Protein expression was Z-score normalized, and hierarchical clustering performed using Ward’s D2. Samples were annotated by experimental groups without clustering. Heatmaps were created using pheatmap. The final processed data table is provided in Table S3.

### Single nucleus RNA sequencing data and visualization

Violin and dot plots were generated using published single-nucleus RNA sequencing (snRNA-seq) data from innervated control muscles, including the tibialis anterior (TA) and gastrocnemius (GAS). Only control (non-denervated) nuclei were used to visualize baseline gene expression patterns across different cell types. The dataset was obtained from Ham et al. [12] which profiled snRNA-seq from mouse skeletal muscle under both control and denervated conditions.

### Statistics

All values were presented as mean=±=S.E.M. with individual animal data points shown in graphs (the number of the points represents n). Data were analyzed in GraphPad Prism 8 and student’s t tests were applied for pairwise comparisons.

## Supporting information

Supplementary Table_1_microdissected_GAS_Muscle_MTJ

Supplementary Table_2_microdissected_TRI_Muscle_MTJ

Supplementary Table_3_average and interaction contrast

Supplementary Table_4_dissected_GAS_Muscle_Tendon

Supplementary Table_5_Antibodies and Probes

Supplementary Code_1_Generation sce

Supplementary Code_2_Differential Expression Analysis

## Acknowledgments

We thank the Biozentrum core facilities for their technical support with imaging (Imaging Core Facility), proteomics (Proteomics Facility), ultrastructural analysis (BioEM Lab), and animal housing (Animal Facility). In particular, we are grateful to the Proteomics Facility and Cinzia Tibera from the BioEM Lab for their expert assistance. We also thank the Bioinformatics Core Facility of the DBM (Department of Biomedicine) for their support in proteomics analysis. Finally, we thank Dr. Judith Reinhard and Dr. Timothy McGowan for their advice on the manuscript. This work was funded by the Swiss National Science Foundation (#189248, #187415, #220245), and the cantons of Basel-Stadt and Basel-Landschaft awarded to M.A.R.

## Supplementary Material

**Supplementary Figure 1:**
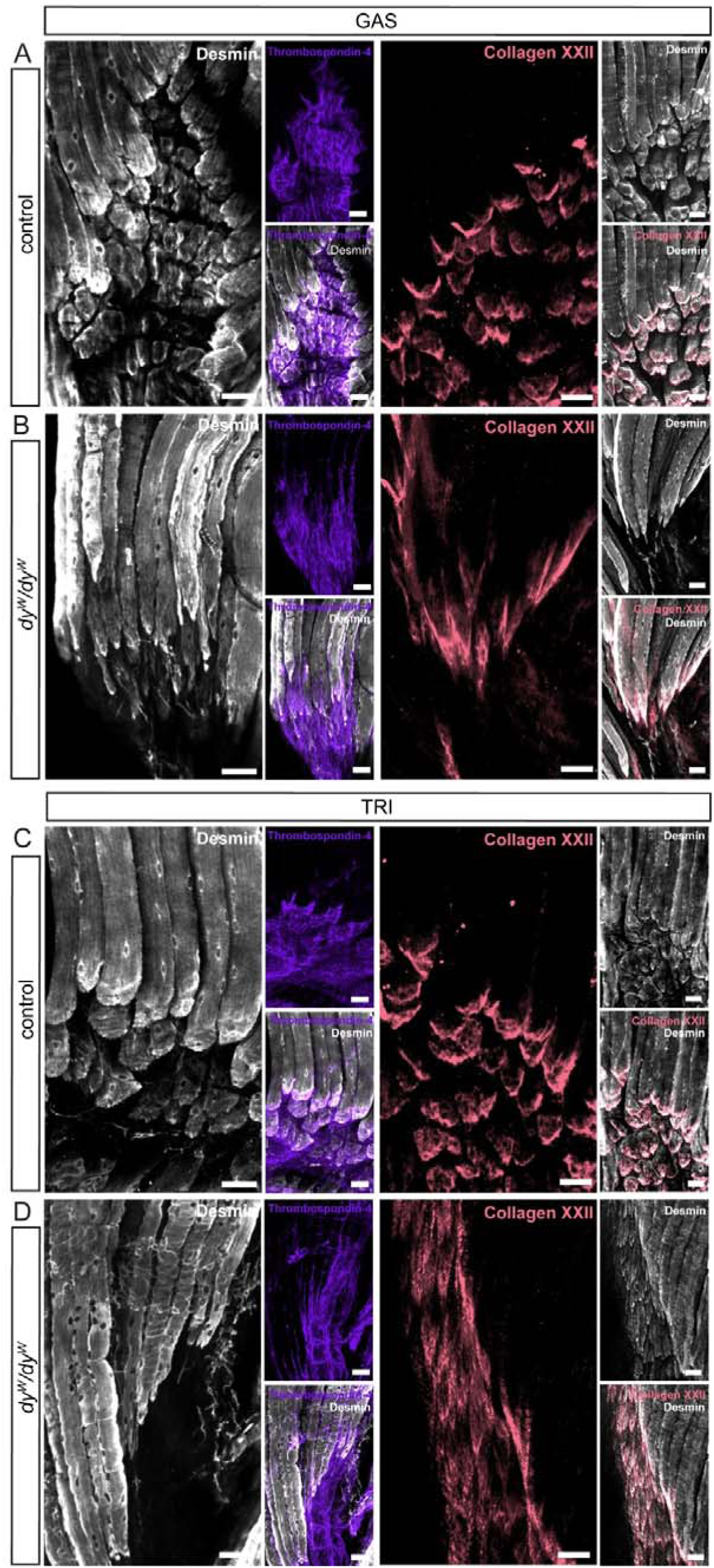
**(A-D)** Confocal micrographs of muscle-tendon whole-mounts from *gastrocnemius* (GAS) (A-B) and *triceps brachii* (TRI) (C-D) from control and *dy^W^/dy^W^* mice. Muscle fibers stained with Desmin (white) insert into the tendon visualized using Thrombospondin-4 (lilac). Muscle fiber tips are covered by Collagen XXII (red). Representative images from n = 5 biological replicates; scale bar = 50 µm.

**Supplementary Figure 2:**
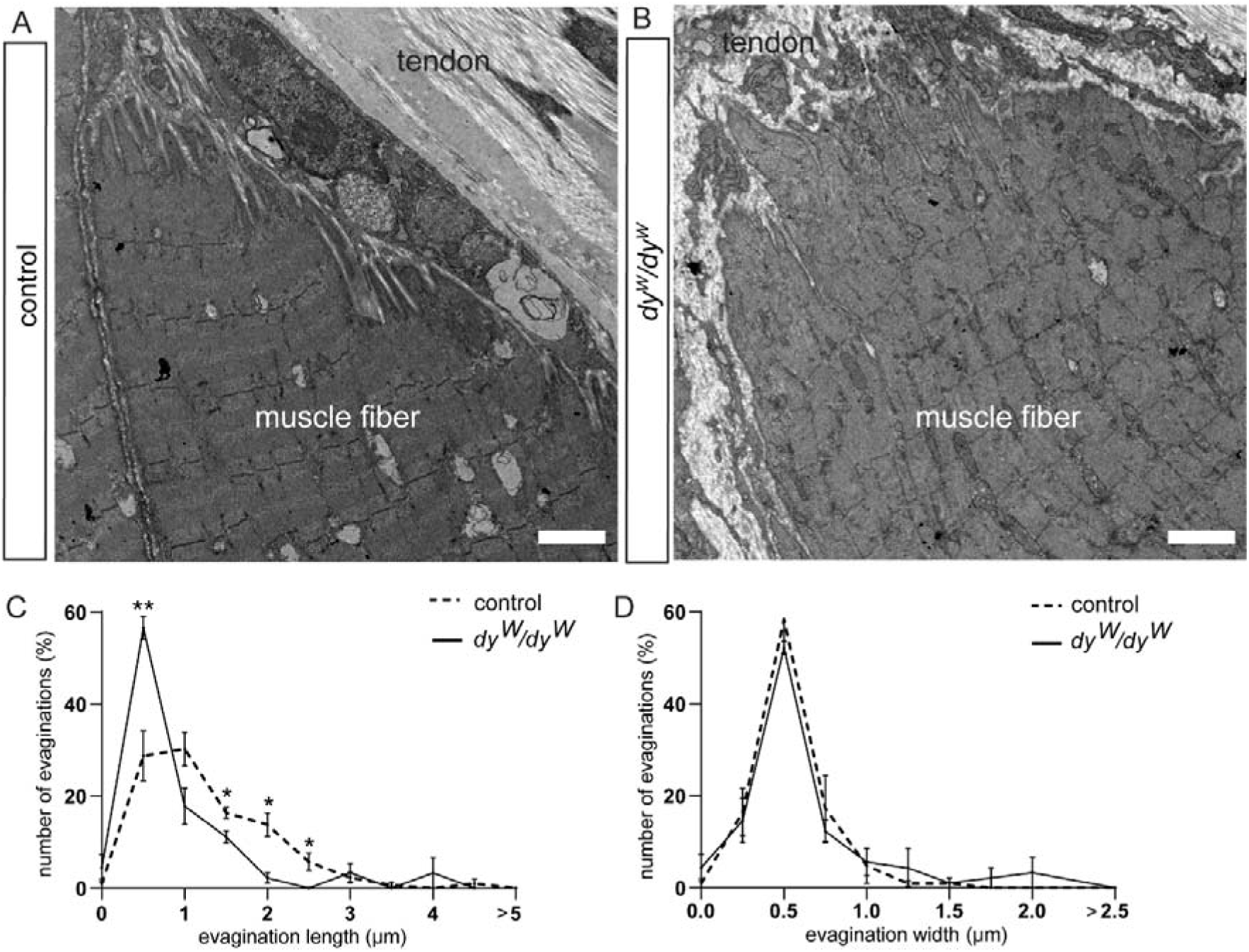
**(A-B)** Transmission-electron micrographs at the muscle-tendon interface of *tibialis anterior* (TA) from control and *dy^W^/dy^W^* mice. **(C-D)** Quantification of the evagination length (C) and evagination width (D) in the TA displayed by the finger-like projections observed in (A-B). Representative images from n = 3 biological replicates; scale bar = 2 µm.

**Supplementary Figure 3:**
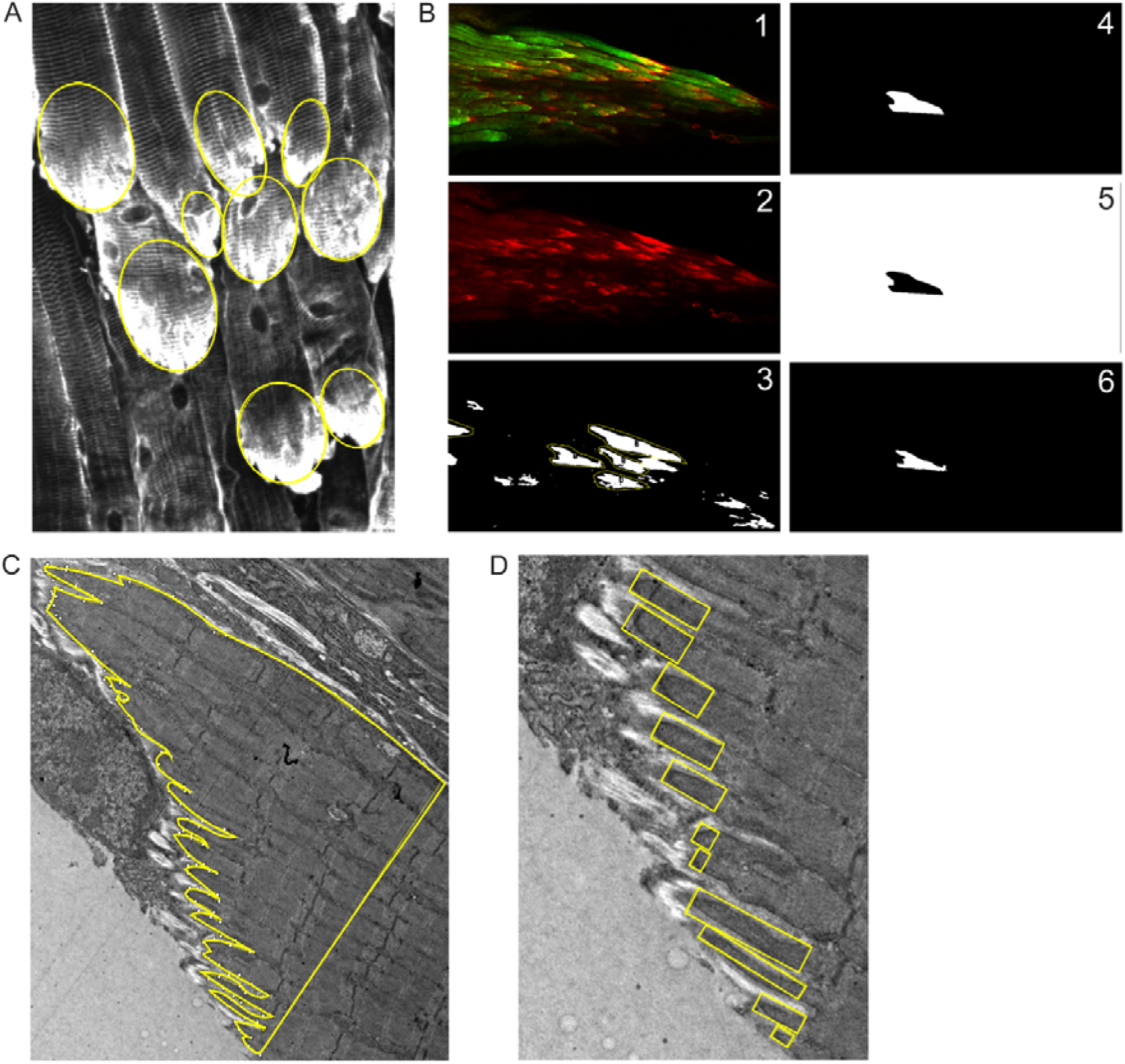
**(A)** Visualization of ROIs helping to measure muscle fiber tip circularity in FIJI. **(B)** Measurement process of the max. Feret diameter of the Collagen XXII staining in FIJI: (1.) composite image, (2.) z projection of Collagen XXII signal, (3.) binary image of the Collagen XXII signal and ROIs, (4.) mask of the Collagen XXII signal of one muscle fiber tip (5.) inversion of the mask (6.) result of the subtraction of the binary image from the mask (7.) measurement of max. Feret diameter (not shown). **(C)** Helping line surrounding the sarcoplasmic evaginations, used to measure the junction length based on the perimeter and diameter of the muscle fiber tip. **(D)** Rectangular ROIs as helping tools for the measurement of sarcoplasmic evagination length and width in FIJI.

**Supplementary Figure 4:**
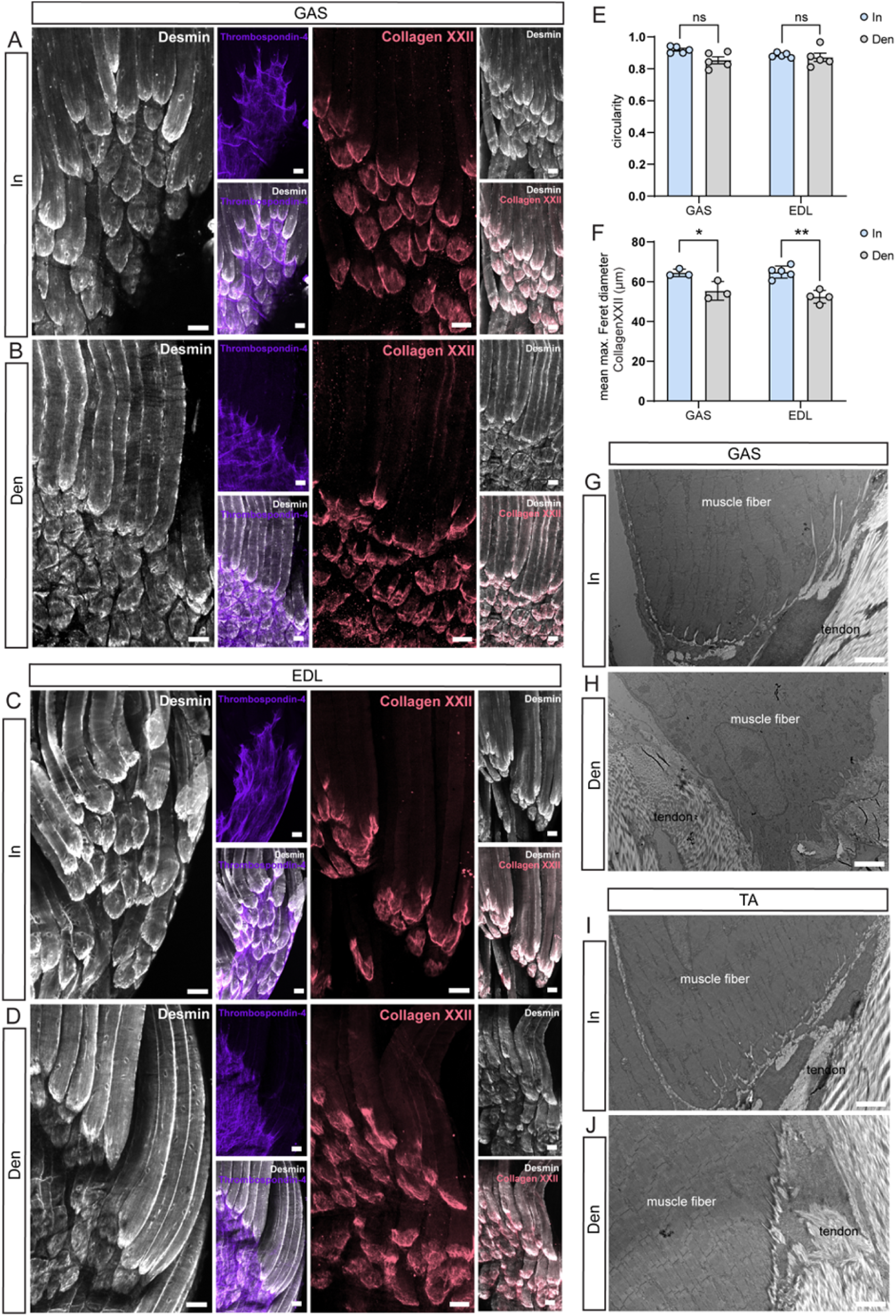
**(A-D)** Confocal micrographs of muscle-tendon whole-mounts from *gastrocnemius* (GAS) (A-B) and *extensor digitorum longus* (EDL) (C-D) from innervated and denervated mice after 1-week denervation. Muscle fibers stained with desmin (white) insert into the tendon visualized using thrombospondin-4 (lilac). Muscle fiber tips are covered by collagen XXII (red). Representative images from n = 5 biological replicates; scale bar = 50 µm. **(E)** Quantification of muscle fiber tip circularity based on desmin immunostaining in (A-D) (n = 5, ns > 0.05; student t-test; mean ± S.E.M.). **(F)** Quantification of the max. Feret diameter of the collagen XXII signal (n = 5, **P < 0.01, *P < 0.05; student t-test; mean ± S.E.M.) **(G-J)** Transmission-electron micrographs at the muscle-tendon interface of innervated and denervated mice from GAS (G-H) and *tibialis anterior* (TA) (I-J). Representative images from n = 3 biological replicates; scale bar = 2 µm.

**Supplementary Figure 5:**
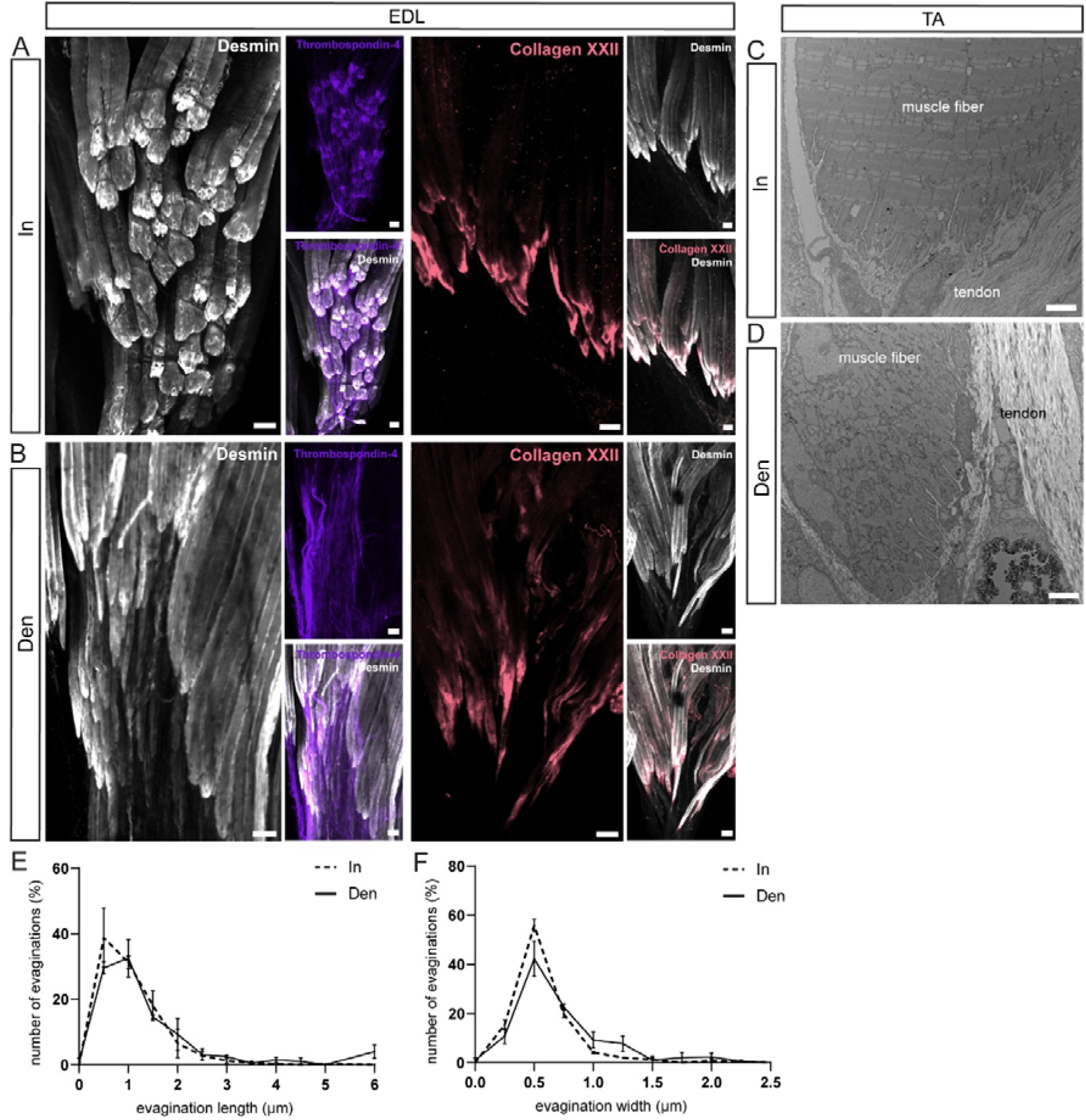
**(A-B)** Confocal micrographs of muscle-tendon whole-mounts from *extensor digitorum longus* (EDL) from innervated and denervated mice after 4 weeks of denervation. Muscle fibers stained with desmin (white) insert into the tendon visualized using thrombospondin-4 (lilac). Muscle fiber tips are covered by collagen XXII (red). Representative images from n = 5 biological replicates; scale bar = 50 µm. **(C-D)** Transmission-electron micrographs of the muscle-tendon interface from *tibialis anterior* (TA) of 4-weeks innervated (C) and denervated (D) mice (n = 3, scale bar = 2 µm). **(E-F)** Quantification of the evagination length (E) and evagination width (F) shown in C-D.

**Supplementary Figure 6:**
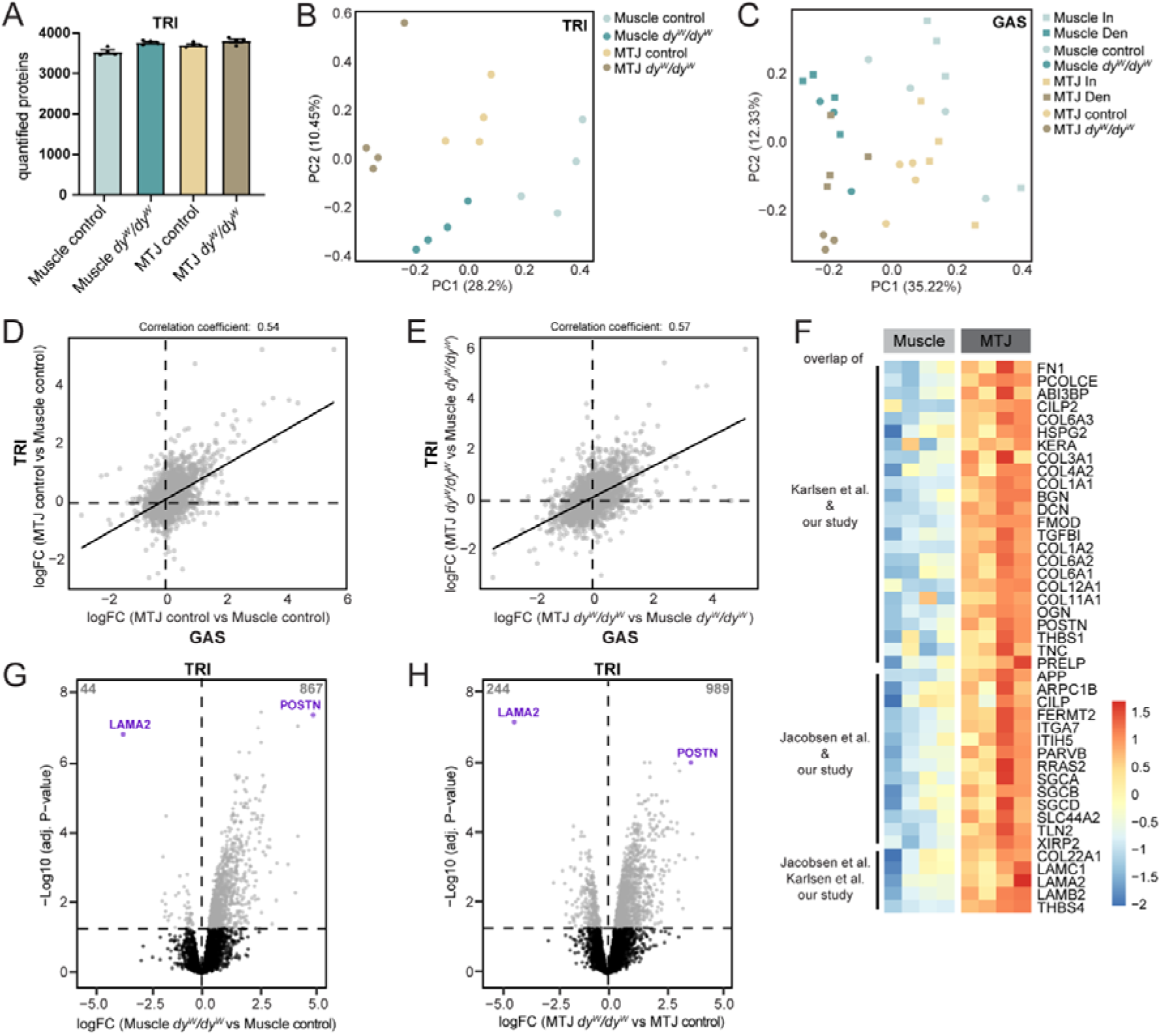
**(A)** Bar plot of the number of quantified proteins within each sample type for TRI proteomics analysis. **(B)** Principal component analysis for TRI showing the segregation of the samples. **(C)** Principal component analysis for GAS showing the segregation of all samples. **(D-E)** Pairwise comparisons of MTJ vs Muscle region within control (D) and *dy^W^/dy^W^*(E) mice of TRI-specific (y-axis) and GAS-specific (x-axis) abundance changes. **(F)** Heatmap showing the relative abundance of MTJ-enriched proteins in control muscle and MTJ samples, which were overlapping between our dataset and those reported previously [6,13]. Proteins are ordered by overlap category, determined in Fig. 4E. Color in each tile represents the scaled abundance value. **(G-H)** Volcano plots displaying the comparison of *dy^W^/dy^W^* vs control proteome in the sample region muscle (G) and MTJ (H) separately. The significance level of the adjusted P-value of 0.05 is indicated by a horizontal dashed line.

**Supplementary Figure 7:**
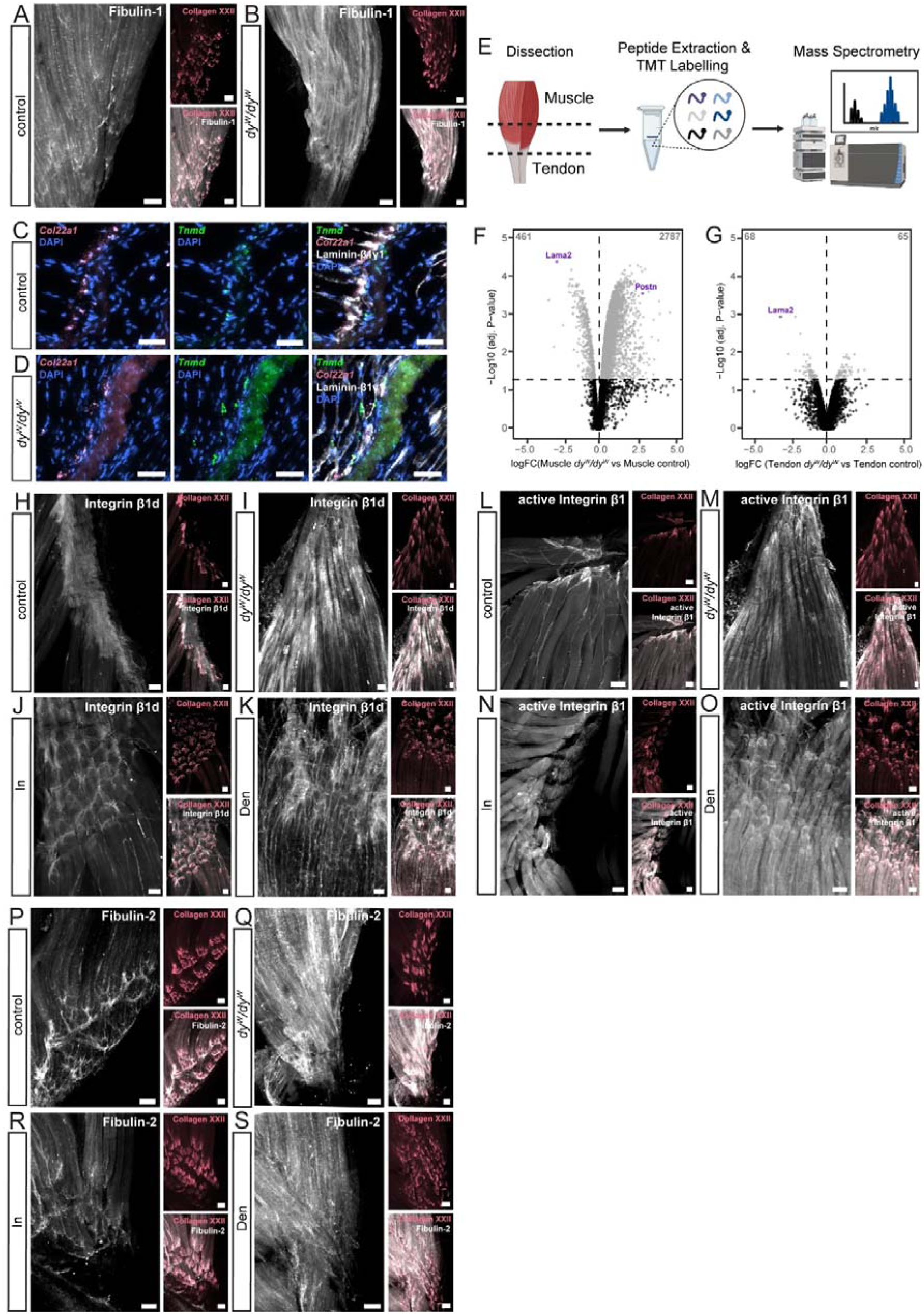
**(A-B)** Confocal micrographs of muscle-tendon whole-mounts stained for Fibulin-1. **(C-D)** Representative images from sm-FISH for *Col22a1* and *Tnmd* on skeletal muscle longitudinal sections stained with laminin-β1-γ1. **(E)** Illustration of the proteomics workflow including dissection of the different sample regions of the GAS muscle-tendon group, sample preparation with TMT labelling and LC-MS. **(F-G)** Volcano plots displaying the comparison of *dy^W^/dy^W^* vs control proteome in the sample region Muscle (F) and Tendon (G) separately. The significance level of the adjusted P-value of 0.05 is indicated by a horizontal dashed line. **(H-K)** Confocal micrographs of muscle-tendon whole-mounts stained with Integrin β1d. **(L-O)** Confocal micrographs of muscle-tendon whole-mounts stained with active Integrin β1. **(P-S)** Confocal micrographs of muscle-tendon whole-mounts stained with Fibulin-2. Representative images from n = 3 biological replicates; scale bar = 50 µm.

