## Supplementary Code_1_Generation sce for "Laminin-α2 is required for the maintenance of the myotendinous junction in vivo": Supplementary Code_1_Generation sce.R

```
# compile rda objects into sce for differential expression analysis
# load important libraries
library(tidyverse)
library(SingleCellExperiment)

# construct SCE object
# load protein summary from dirpath <- path to protein_summary

load(file.path(dirpath, "protein_summary.rda"))
load("protein_summary.rda")

# counts
Log2Int <- as.matrix(protein_summary[, grep("Log2Int_", colnames(protein_summary))])
rownames(Log2Int) <- protein_summary$GeneName

# rowData (remaining columns of protein_summary)
rd <- as.data.frame(protein_summary[, grep("Log2Int_", colnames(protein_summary), invert = TRUE)])
rownames(rd) <- rd$GeneName

# colData (sample information)
library(stringr)
cd <- data.frame(combined = colnames(protein_summary)[grep("Log2Int_", colnames(protein_summary))])

# cd <- cd %>% separate_wider_delim(combined, delim = "-", names = c("prefix", "suffix"))
# cd$number <- str_extract(cd$prefix, "[0-9]{3}")
# load additional metadata
metadata <- read.table("metadata_all.csv",
                       header = TRUE, sep = ";")

cd <- full_join(cd, metadata, by = join_by(combined == sample.name))

cd$plateLoc <- str_extract(cd$combined, "[A-Z]{1}[0-9]{1}")

cd <- dplyr::rename(cd, sample.name = combined)
cd$condition <- sapply(cd$group, function(x) strsplit(x, "_")[[1]][2])

# columns as factors
# and relocate columns
cd <- mutate(cd, group = factor(group), 
             region = factor(region), condition = factor(condition),
             mouse.number = factor(mouse.number), plateLoc = factor(plateLoc)) %>%
  relocate(sample.name, group, region, condition, mouse.number, plateLoc)

sce <- SingleCellExperiment(assays = list(logcounts = Log2Int),
                            colData = cd,
                            rowData = rd)

colnames(sce) <- make.names(colnames(sce))
saveRDS(sce, "sce.rds")

# remove outliers: two KO samples 
# (based on Lama2 expression and clustering in heatmap to WT)
outliers <- c("Log2Int_007_LMDlow6_DIA_Y.G1_2190_H24", "Log2Int_015_LMDlow6_DIA_Y.G2_2186_H24")
colData(sce)[outliers, ]
# mouse.number 40471 affected

sce <- sce[, !colnames(sce) %in% outliers]
sce$mouse.number <- droplevels(sce$mouse.number)
saveRDS(sce, "sce.rds")
```
