## Supplementary Code_2_Differential Expression Analysis for "Laminin-α2 is required for the maintenance of the myotendinous junction in vivo": Supplementary Code_2_Differential Expression Analysis.R

```
# Differential expression analysis was done on the sce object generated from the proteomics data file

# Question 1: KO vs WT

# relevel to Muscle_WT
sce$condition <- relevel(sce$condition, ref = "WT")
sce$region <- relevel(sce$region, ref = "Muscle")

# link between mouse.number and treatments
# condition is nested inside mouse.number
# table(sce$mouse.number, sce$condition)
table(sce$mouse.number)
table(sce$condition, sce$region, sce$mouse.number)

design_WTKO_paired <- model.matrix(~ 0 + mouse.number, data = colData(sce))

# paired samples (same mouse.number)
WT_MTJ_vs_Muscle <- sce$condition == "WT" & sce$region == "MTJ"
KO_MTJ_vs_Muscle <- sce$condition == "KO" & sce$region == "MTJ"
design_WTKO_paired <- cbind(design_WTKO_paired,
                WT_MTJ_vs_Muscle,
                KO_MTJ_vs_Muscle)

design_WTKO_paired

# Fit linear model for paired data
fit_WTKO_paired <- lmFit(logcounts(sce), design_WTKO_paired)

# Define contrasts for paired comparisons
contrasts_WTKO_paired <- makeContrasts(
  WT.MTJvsMuscle = WT_MTJ_vs_Muscle,
  KO.MTJvsMuscle = KO_MTJ_vs_Muscle,
  Interaction.MTJvsMuscle = KO_MTJ_vs_Muscle - WT_MTJ_vs_Muscle,
  Average.MTJvsMuscle = (KO_MTJ_vs_Muscle + WT_MTJ_vs_Muscle) / 2,
  levels = colnames(design_WTKO_paired)
)

fit3_WTKO_paired <- contrasts.fit(fit_WTKO_paired, contrasts_WTKO_paired)
fit3_WTKO_paired <- eBayes(fit3_WTKO_paired)

WT_MTJvsMuscle <- topTable(fit3_WTKO_paired, coef = "WT.MTJvsMuscle", number ="Inf")
KO_MTJvsMuscle <- topTable(fit3_WTKO_paired, coef = "KO.MTJvsMuscle", number ="Inf")

interactionWTKO <- topTable(fit3_WTKO_paired, coef = "Interaction.MTJvsMuscle", number ="Inf")
averageWTKO <- topTable(fit3_WTKO_paired, coef = "Average.MTJvsMuscle", number ="Inf")

#unpaired (addition)
design_WTKO_unpaired <- model.matrix(~ 0 + condition:region, data = colData(sce))

colnames(design_WTKO_unpaired) <- make.names(colnames(design_WTKO_unpaired))  # Clean column names

# Fit linear model
fit_WTKO_unpaired <- lmFit(logcounts(sce), design_WTKO_unpaired)

contrasts_WTKO_unpaired <- makeContrasts(
  KO.MTJvsWT.MTJ = conditionKO.regionMTJ - conditionWT.regionMTJ,
  KO.MusclevsWT.Muscle = conditionKO.regionMuscle - conditionWT.regionMuscle,
  levels = colnames(design_WTKO_unpaired)
)

fit3_WTKO_unpaired <- contrasts.fit(fit_WTKO_unpaired, contrasts_WTKO_unpaired)
fit3_WTKO_unpaired <- eBayes(fit3_WTKO_unpaired)

KO_MTJ_vs_WT_MTJ <- topTable(fit3_WTKO_unpaired, coef = "KO.MTJvsWT.MTJ", number = "Inf")
KO_Muscle_vs_WT_Muscle <- topTable(fit3_WTKO_unpaired, coef = "KO.MusclevsWT.Muscle", number = "Inf")


# Question 2: DEN vs IN

# relevel to Muscle_IN
sce$condition <- relevel(sce$condition, ref = "IN")
sce$region <- relevel(sce$region, ref = "Muscle")

# link between mouse.number and treatments
# condition is nested inside mouse.number
# table(sce$mouse.number, sce$condition)
table(sce$mouse.number)
table(sce$condition, sce$region, sce$mouse.number)

design_DENIN_paired <- model.matrix(~ 0 + mouse.number, data = colData(sce))

# paired samples (same mouse.number)
IN_MTJ_vs_Muscle <- sce$condition == "IN" & sce$region == "MTJ"
DEN_MTJ_vs_Muscle <- sce$condition == "DEN" & sce$region == "MTJ"

design_DENIN_paired <- cbind(design_DENIN_paired,
                  IN_MTJ_vs_Muscle,
                  DEN_MTJ_vs_Muscle)
design_DENIN_paired

# Fit linear model
fit_DENIN_paired <- lmFit(object = logcounts(sce), design = design_DENIN_paired,
               method = "ls")

contrasts_DENIN_paired <- makeContrasts(
  IN.MTJvsMuscle = IN_MTJ_vs_Muscle,
  DEN.MTJvsMuscle = DEN_MTJ_vs_Muscle,
  
  Interaction.DENvsIN = DEN_MTJ_vs_Muscle - IN_MTJ_vs_Muscle,
  Average.DENvsIN = (DEN_MTJ_vs_Muscle + IN_MTJ_vs_Muscle) / 2,
  
  levels = colnames(design_DENIN_paired))

contrasts_DENIN_paired

fit3_DENIN_paired <- contrasts.fit(fit_DENIN_paired, contrasts_DENIN_paired)
fit3_DENIN_paired <- eBayes(fit3_DENIN_paired)

IN_MTJvsMuscle <- topTable(fit3_DENIN_paired, coef = "IN.MTJvsMuscle", number = "Inf")
DEN_MTJvsMuscle <- topTable(fit3_DENIN_paired, coef = "DEN.MTJvsMuscle", number = "Inf")

interactionDENIN <- topTable(fit3_DENIN_paired, coef = "Interaction.DENvsIN", number = "Inf")
averageDENIN <- topTable(fit3_DENIN_paired, coef = "Average.DENvsIN", number = "Inf")


#unpaired (Zusatz)
design_DENIN_unpaired <- model.matrix(~ 0 + condition:region, data = colData(sce))

colnames(design_DENIN_unpaired) <- make.names(colnames(design_DENIN_unpaired))  # Clean column names

# Fit linear model
fit_DENIN_unpaired <- lmFit(logcounts(sce), design_DENIN_unpaired)

contrasts_DENIN_unpaired <- makeContrasts(
  DEN.MTJvsIN.MTJ = conditionDEN.regionMTJ - conditionIN.regionMTJ,
  DEN.MusclevsIN.Muscle = conditionDEN.regionMuscle - conditionIN.regionMuscle,
  levels = colnames(design_DENIN_unpaired)
)

fit3_DENIN_unpaired <- contrasts.fit(fit_DENIN_unpaired, contrasts_DENIN_unpaired)
fit3_DENIN_unpaired <- eBayes(fit3_DENIN_unpaired)

DEN_MTJ_vs_IN_MTJ <- topTable(fit3_DENIN_unpaired, coef = "DEN.MTJvsIN.MTJ", number = "Inf")
DEN_Muscle_vs_IN_Muscle <- topTable(fit3_DENIN_unpaired, coef = "DEN.MusclevsIN.Muscle", number = "Inf")
```
